# The rhythmic transcriptional landscape in *Caenorhabditis elegans*: daily, circadian and novel 16-hour cycling gene expression revealed by RNA-sequencing

**DOI:** 10.1101/2024.07.06.602329

**Authors:** Jack Munns, Katherine Newling, Sally R. James, Lesley Gilbert, Seth J. Davis, Sangeeta Chawla

**Affiliations:** Department of Biology, University of York, York, UK; York Biomedical Research Institute, University of York, York, UK

## Abstract

The nematode *Caenorhabditis elegans* is an unconventional model in chronobiology, reported to exhibit physiological and behavioural circadian rhythms while lacking a defined transcriptional oscillator. The extent and importance of circadian rhythms in *C. elegans* remain uncertain, given a probable lack of functional conservation of key circadian proteins, relatively non-robust reported rhythms and an ambiguous diel ecology. Here, we investigated the temporal coordination of gene expression in *C. elegans* post-development using RNA sequencing. Over a circadian time series, in which we synchronised nematodes to combined light and temperature cycles, we found clear evidence of daily oscillations in 343 genes using JTK_Cycle. However, rhythms were not well-sustained in constant conditions, with only 13 genes remaining significantly rhythmic. Reanalysis of previous transcriptomic data echoed this finding in identifying far fewer rhythmic genes in constant conditions, while also identifying a greater number of rhythmic genes overall. Weighted gene co-expression network analysis (WGCNA), a hierarchical clustering approach, further confirmed prevalent environmentally driven daily oscillations in the RNA-seq data. This analysis additionally revealed a novel co-expression trend in which over 1000 genes exhibited hitherto unreported 16-hour oscillations, highlighting a new facet of temporal gene expression coordination in *C. elegans*.

## Introduction

Biological oscillations enable temporal coordination of cellular and organismal biology. Circadian rhythms are an evolutionarily pervasive example of such oscillations, with approximately 24-hour physiological rhythms being widely observable in animals, fungi, plants and photosynthetic prokaryotes and having been reported in non-photosynthetic prokaryotes [1–5]. The physiological processes that exhibit 24-hour rhythms are diverse, but common to circadian model organisms, and potentially underpinning physiological rhythms, are rhythms in gene expression. In animal models, transcriptomic studies have reported gene expression rhythms to extend to hundreds to thousands of genes in *Drosophila* [6] and over 43% of protein-coding genes in mice [7].

Within the metazoa, part-conserved transcriptional oscillators are proposed to drive rhythms in gene expression and wider rhythmicity. Transcription-translation feedback loop (TTFL) mechanisms centre around circadian oscillations in the RNA and protein abundance of homologues of the *period* (*per*) gene, first identified in *Drosophila* [8–10]. The *per gene* is conserved in mammals (which express three *Per* genes) and other vertebrates [11,12]. In the respective TTFL models, *per*/*Per* transcription is driven by similarly homologous heterodimeric basic helix-loop-helix (bHLH) transcription factors which the resulting PER homologue protein products interact with and repress as part of autorepressor complexes. Regulated degradation of the repressor complexes allows transcription to resume, resulting in an endogenously generated, delayed feedback loop. The approximate 24-hour periodicity of the resulting autorepressor loops is proposed to result in cell autonomous circadian rhythms in both the repression of the respective transcription factors and downstream gene expression [1,13].

The nematode *Caenorhabditis elegans* is a curious case among laboratory model organisms in that it has been reported to exhibit numerous behavioural and physiological circadian rhythms but has no described TTFL. Reported circadian rhythms in *C. elegans* include gene expression [14–16], locomotor activity [17–22], protein abundance, oxidation-reduction cycles, olfactory responses [15,23], feeding, oxygen consumption [24], enzymatic activity [25] and hyperosmotic stress resistance [26]. Some of these reported rhythms do appear to be relatively non-robust however, being detectable at the level of select populations or varying in their activity peaks [16,18–20]. Phylogenetically, *C. elegans* is positioned in the Ecdysozoa within the protostomes [30,31] and as such, a common ancestral basis of rhythmicity shared between insects and mammals would plausibly be shared with *C. elegans*. However, the *C. elegans* sequence homologue of *per*, *lin-42*, and other identified sequence homologues of oscillatory TTFL genes [27,28], have been reported not to oscillate in adult nematodes [14,15,29]. Many genes, including *lin-42*, do oscillate throughout *C. elegans*’ development, however [32–40]. *lin-42* in particular is well-described as a heterochronic gene necessary for progression through a temporally defined series of larval stages [41–47], highlighting an alternate function of this key TTFL homologue and biological timing more broadly in nematodes.

*C. elegans* also notably differs from the more established *Drosophila* and mammalian circadian models with respect to its evolutionary ecology. Light is the prevailing synchronising cue used in the study of circadian rhythms in both invertebrate and vertebrate chronobiology models [1,11], and *C. elegans* is photoreceptive and behaviourally phototactic [48,49]. However, only one directly light-sensitive photoreceptor protein has been identified in *C. elegans*, the UV-sensitive LITE-1 [50–52]. Nematodes also lack the evolutionarily conserved cryptochrome (CRY) proteins [28,53], key in transducing photic inputs to the *Drosophila* circadian clock [54] and for circadian transcriptional repression in mammals [55,56]. *C. elegans*’ natural ecology (and thus its exposure to daily environmental cycles) is also not well-understood, with populations having been isolated from various surface level and subterranean environments [57–61]. It exhibits a boom-and-bust life cycle, including periods of fast population growth and inactivity; individuals are frequently isolated as dauers [58], a metabolically less active, stress-resistant and diapause state entered into when population density, food availability or temperature conditions become unfavourable [62,63]. The extent to which *C. elegans* has evolved to interact with a 24-hour diel environment is therefore unclear.

*C. elegans* is therefore an atypical model of biological timing, reported to show numerous physiological circadian rhythms, but potentially lacking the functional conservation of the transcriptional oscillators seen in *Drosophila* and mammals, while also differing in its diel ecology. As such, questions remain as to the importance and nature of its circadian system. Here, we employed transcriptomics to comprehensively investigate gene expression oscillations in *C. elegans* within a circadian context, post-development. To date, very few circadian transcriptomic studies have been performed in *C. elegans*, but hundreds of endogenous cycling genes synchronised by light or temperature cycles have been reported [14,29]. We build upon prior work in conducting an RNA sequencing time series in *C. elegans* using a sextuplicate sampling approach. In so doing, we identify approximately 24-hour rhythms in gene expression, but not to the same extent as a prior dataset [14], including upon reanalysis here. Rather than endogenous circadian rhythms, we identify a greater prevalence of genes that show variation in line with environmental cycles. Finally, we also identify a large subset of phase-coherent genes with approximately 16-hour ultradian periodicity that are, to our knowledge, previously unreported in nematodes not undergoing development.

## Results

### Sampling *C. elegans* messenger RNA over a circadian time series

To investigate circadian gene expression rhythms in *C. elegans*, we used combined 12:12-hour cycles of light and temperature to synchronise (entrain) nematodes (light at 20°C: dark at 15°C). Our rationale for using these dual environmental inputs was that *C. elegans* is sensitive to both light [49,50] and temperature [64,65] and both sensory inputs have previously been reported to function as entrainment cues (*zeitgebers*; ‘time givers’) for gene expression rhythms in *C. elegans* [14–16,29,66]. While temperature has been used to entrain rhythms in other more established poikilothermic models, light is the better studied and reportedly more effective *zeitgeber* [1,11,67–71]. We therefore reasoned that using both light and temperature would be a sensible approach to detect rhythmic genes. We entrained nematodes to combined light and temperature cycles from embryogenesis and for three days as adults before releasing the animals into constant darkness at 15°C.

For our sampling approach, we considered that *C. elegans* rhythms could be relatively non-robust compared to conventional chronobiology models (with certain rhythms being reportedly detectable at population level in some populations, for example [16,18,19]). To maximise rhythmic gene detection while incorporating endogenous timing, sampling of mRNA was performed across the final entrainment day and first day of constant conditions at 4-hour intervals. We also considered that conventional triplicate sampling could miss modest expression changes in a relatively non-robust system [72]. As such, to minimise false negative results and capture the extent of differential gene expression as accurately as possible, we sampled in sextuplicate; we harvested mRNA from 6 replicate populations of nematodes per timepoint (with one replicate being excluded from analysis at the final timepoint due to very few reads being detected). Each population comprised approximately 150 age-synchronised adult hermaphrodites housed on a petri plate (Figure 1).

**Figure 1:**
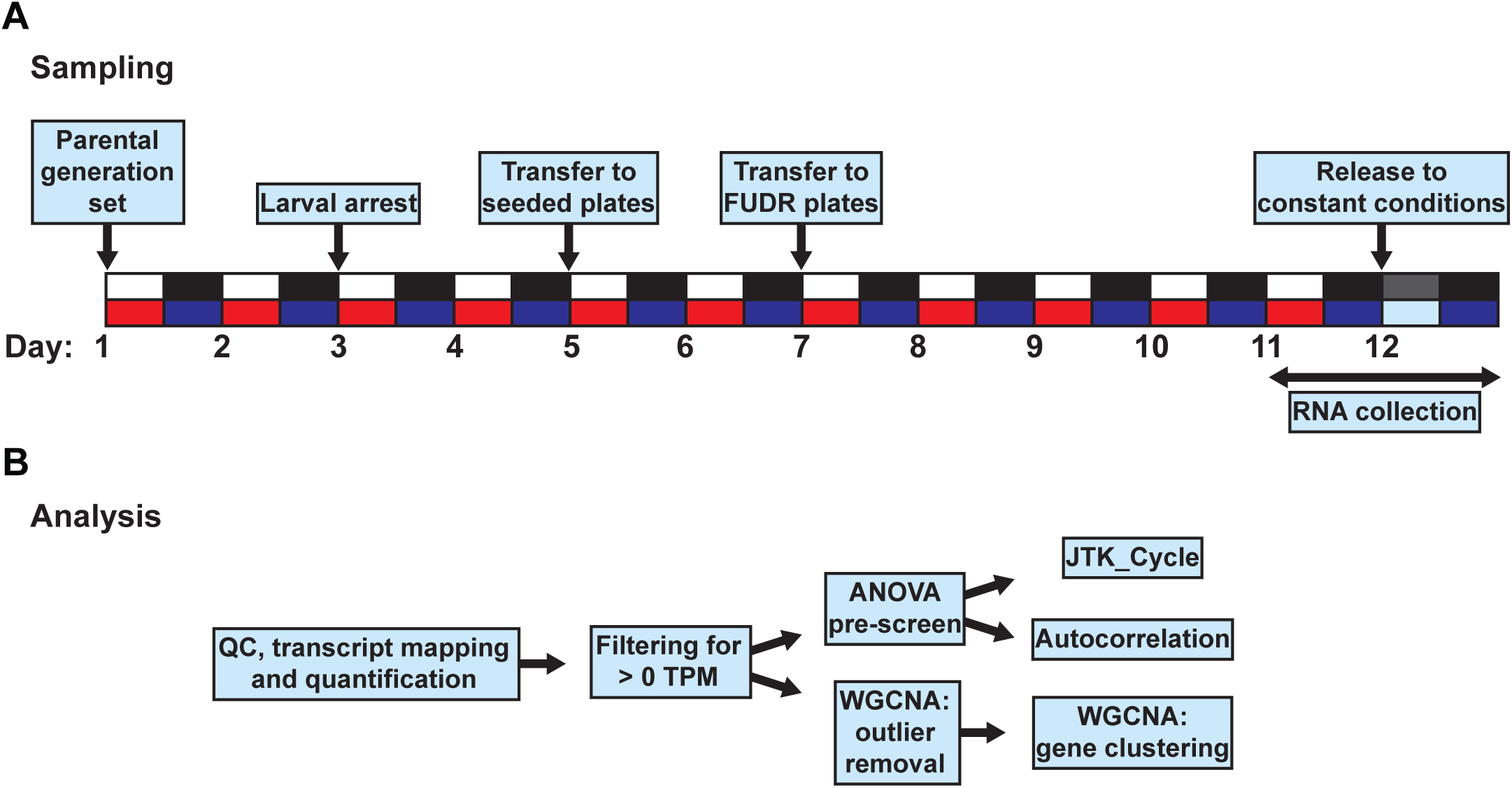
RNA-seq (A) Entrainment and Sampling Protocol and (B) Analysis. **A:** Nematodes were exposed to 12:12-hour cycles of light at 20°C (white and red boxes) and dark at 15°C (black and dark blue boxes). Parental individuals were entrained until gravid and eggs were isolated by bleaching (see Methods). Offspring were maintained in starvation conditions to generate age-synchronised individuals before being moved to plates seeded with *Escherichia coli*. At the L4 larval stage, nematodes were transferred to plates also containing 25 μM 5-fluorodeoxyuridine (FUDR) to prevent reproduction. Nematodes were entrained for 3 days as adults and then released into constant dark at 15°C. RNA harvesting was performed every 4 hours on the final day of entrainment and first day in constant conditions (day 10 and day 11) with each replicate being a *C. elegans* population from one petri plate. Following sequencing and mapping (described in Methods), we filtered transcripts to include those with > 0 transcripts per million (TPM) and employed two analysis approaches: first, an ANOVA pre-screen followed by JTK_Cycle [73,74] and autocorrelation analysis to identify circadian rhythms; second, weighted gene co-expression network analysis (WGCNA) [75] to identify expression trends without prior assumptions as regards to periodicity or waveform.

### JTK_Cycle reveals genes expressed with approximately 24-hour periodicity

To identify genes with circadian expression patterns, we first analysed time-series data using JTK_Cycle [73,74], a widely used method for identifying transcriptome-wide circadian rhythms in diverse model organisms [6,7,76–79]. Genes were first filtered to include only those for which 50% of all samples showed an expression value > 0 (in transcripts per million, TPM; Figure 1B). From this initial list of 16,716 genes, we applied an ANOVA pre-screen (*p* < 0.05) to identify genes that showed significant differences in TPM between timepoints. On the resulting set of 2526 differentially expressed genes by ANOVA, JTK_Cycle identified 343 significantly rhythmic genes over the full two-day time series (Benjamini-Hochberg-adjusted *p*-value, BHQ < 0.05; full list in Table S1). Genes with the lowest BHQ values included examples of repeating 24-hour oscillations, as well as genes with 28-hour periods that part-replicated the environmental cycle in the absence of external stimuli (Figure 2A). The genes identified by this approach notably varied by phase with peaks or troughs occurring across the circadian cycle (Figure 2B).

**Figure 2:**
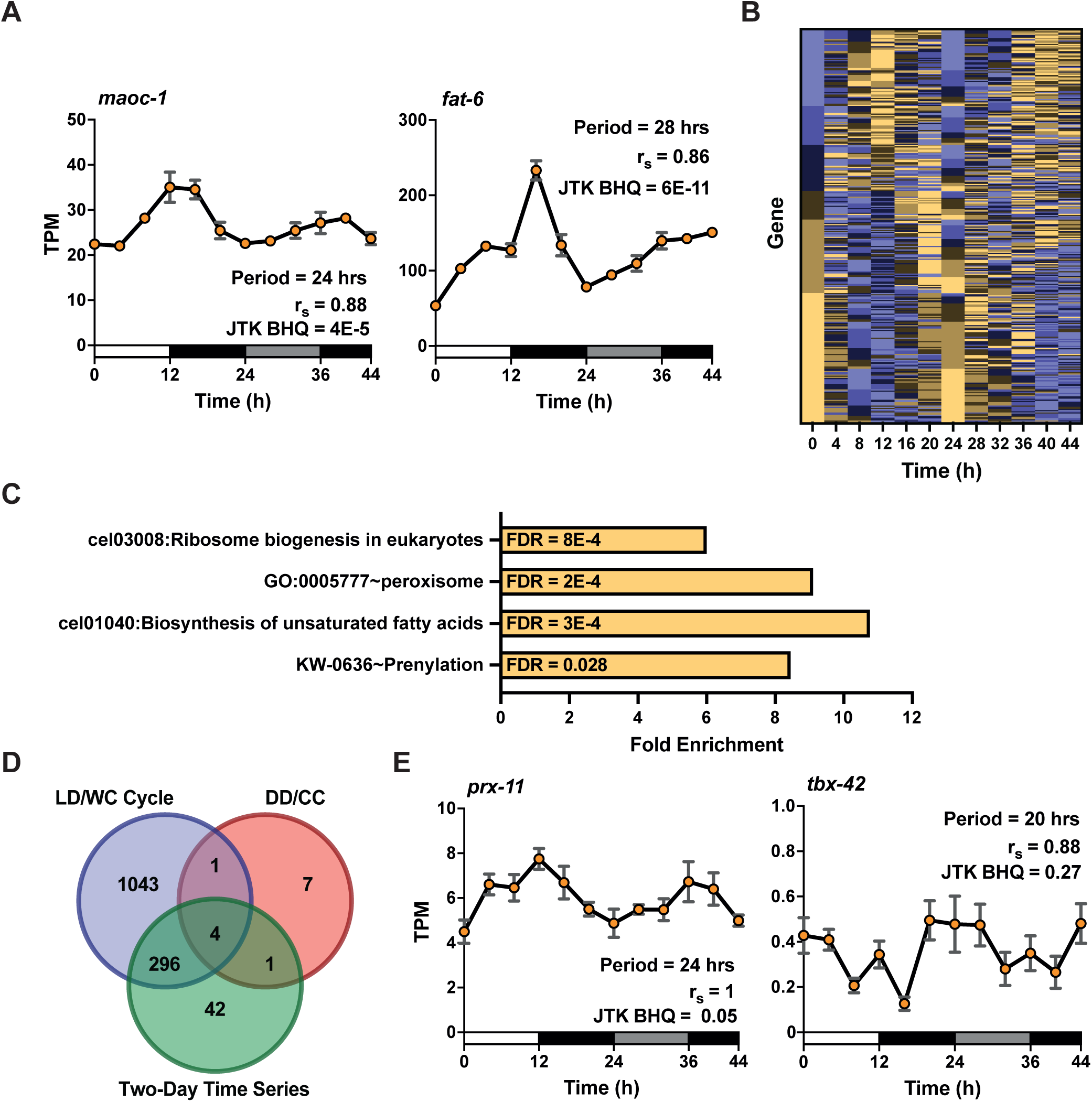
Rhythmicity analysis of a two-day RNA-seq time series. **A:** Examples of genes with 24 and 28-hour periodicities as identified by JTK_Cycle over the full two-day time series, including JTK_Cycle period estimate, Spearman’s rank coefficient, and JTK_Cycle BHQ values (both are within the lowest 7 JTK_Cycle BHQ values). Shown are average TPM values of 6 replicates ± SEM. Time 0 represents the time of simultaneous light onset and temperature increases with environmental conditions indicated by boxes along the *x*-axis (white = light/20°C, black = dark/15°C, light grey = subjective day (dark/15°C)). **B:** Heat map illustrating acrophases and bathyphases of all 343 JTK_Cycle, BHQ < 0.05 genes with brown representing peaks and blue representing troughs. **C:** DAVID enrichment analysis of genes in **B**. Fold enrichment is shown for select DAVID annotation terms with FDR < 0.05 (from four enrichment clusters containing significant terms; only the most significant term in each cluster is shown here). **D:** Venn diagram showing the number of genes collectively identified by JTK_Cycle (BHQ < 0.05) under a 12:12-hour environmental cycle, constant conditions and over the full two-day time series. **E:** Examples of genes with high 24-hour *r_s_* autocorrelation, both identified and not identified as rhythmic by JTK_Cycle (BHQ < 0.05) over the full two-day time series.

To investigate the biological processes potentially served by these rhythmic genes, we performed functional annotation analysis on the 343 JTK_Cycle-significant genes using DAVID (Database for Annotation, Visualization and Integrated Discovery) [80,81]. DAVID compiles annotation terms from multiple sources and generates enrichment clusters of similar terms. Four enrichment clusters contained at least one significant term in our analysis (FDR threshold < 0.05), highlighting ribosome biogenesis, peroxisome function, fatty acid synthesis and prenylation as biological functions associated with our rhythmic gene list (Figure 2C; full annotation cluster results in Table S2).

Notably absent from our list of significantly rhythmic genes were homologues of TTFL-associated genes, consistent with prior literature [14,15,29]. We considered *lin-42* and 10 other genes reported to be sequence homologues of genes associated with TTFLs or circadian clock regulation in *Drosophila* and mammals [27,28,82] (including genes expressed rhythmically and non-rhythmically in these species). While all of these genes met the criterion of detectable expression (TPM > 0 in > 50% of RNA-seq samples), all but two, *atf-2* and *pdfr-1* (related to the *vrille* and PDF receptor genes in *Drosophila* respectively), did not pass our ANOVA pre-screen (*p* > 0.05). We applied JTK_Cycle independently to these 11 genes over our two-day time series and identified no significant circadian rhythmic expression (minimum *p*-value 0.44, in *pdfr-1*; Table S3; Figure S1).

Our experimental design included both an environmental cycle and constant conditions, following the rationale that an entrained endogenous rhythmic waveform should follow an entrained cycle. To test this and assess the relative role of each condition, we also analysed each condition independently, dividing our 12-point time series into two 6-point datasets prior to ANOVA pre-screening. While limited to a maximum period of 24 hours, we could then perform JTK_Cycle on these single-day data. Far more genes showed significant changes in expression over an environmental cycle than constant conditions (2377 and 406 differentially expressed genes respectively; ANOVA *p* < 0.05). Of these, an increased number and proportion were significantly rhythmic over single environmental cycle day (1344 genes with JTK_Cycle BHQ < 0.05, 57% of differentially expressed genes; Table S4) relative to the full two-day time series (343 genes, 15%). Conversely, a much smaller proportion, and very few genes overall, met this statistical threshold in constant conditions (13 genes, 3%; Table S5). While fewer significant rhythms would be expected in constant conditions, consistent with the notion that an entrainment cue serves to synchronise independent, uncoupled cellular oscillators that dampen over time without external inputs [83,84], these results likely suggest a principal role of the light and temperature cycle in generating rhythmic significance over the full two-day analysis.

Of note, only four genes co-occurred between JTK_Cycle analyses; *sws-1*, *C27B7.9*, *gstk-1* and *fat-6* were the only genes identified as significantly rhythmic in an entrainment cycle, constant conditions and the full two-day time series (Figure 2D). We therefore sought to validate our JTK_Cycle results *via* an alternative analysis approach. Because an endogenously regulated circadian gene should exhibit repeating, correlated expression over multiple cycles, we calculated 24-hour rank autocorrelation coefficients for our 2526 differentially expressed genes. Most (233 genes, 67.9%) of the JTK_Cycle-significant genes over the full two-day time series were positively correlated (Spearman’s rank, rs > 0;, 53 met an arbitrary correlation threshold of rs > 0.7 (Table S1, Figure 2A). Notably, however, many genes with high rank correlation coefficients were not previously identified as rhythmic by JTK_Cycle; applying a correlation threshold of rs > 0.7 identified a further 200 highly correlated but previously undetected genes (Figure 2E; Table S1), including one TTFL-associated homologue, *pdfr-1* (Figure S1). Our RNA-seq analyses therefore provide both a small validated subset and an expanded candidate list of genes that exhibit daily cycles for further consideration.

### The rhythmic genes identified in our time series differ in identity from prior time series data

To compare our work to prior findings, we next reanalysed an earlier *C. elegans* transcriptomics dataset (henceforth referred to as ‘the 2010 dataset’; GEO: GSE23528 [14]). The previous study, using microarrays, reported approximately 24-hour expression rhythms under light or temperature cycles and in subsequent constant conditions. As well as applying light and temperature independently, this work used a wider temperature range to entrain *C. elegans* (25:15°C to our 20:15°C). Using a different analysis approach, the previous work reported 692 unique genes to oscillate in cycling and constant conditions, only 15 were reidentified in our RNA-seq JTK_Cycle analysis (BHQ < 0.05, data not shown). We therefore sought to reanalyse this dataset, using an analysis approach comparable to that used for the RNA-seq data. The 2010 dataset consisted of 22,625 transcripts and was similarly generated from 44-hour replicate time series comprising an entrainment day (12:12 hours of light:dark or warm:cold cycles respectively) and a day of constant conditions. This enabled us to perform our analyses (ANOVA pre-screen followed by JTK_Cycle) over combined two-day time series and independent days, as with the RNA-seq data.

This reanalysis echoed our RNA-seq results with respect to finding far more cycling genes when incorporating environmental cycle data compared to constant conditions alone (Figure 3A, C; Table S6). It was also consistent with the published analysis in finding temperature rather than light to be a more powerful driver of oscillations. Surprisingly however, this analysis suggested the extent of rhythmic gene expression to be far greater than the RNA-seq data; far more significant genes were identified under light and particularly temperature entrainment protocols, equating to approximately 25% of transcripts (871 and 4585 genes respectively, BHQ < 0.05). Most genes in these datasets were best identified under a two-day time series, as opposed to the environmental cycle day seen in the RNA-seq data, possibly suggesting more coherence in gene expression between cycling and constant conditions (Figure 3A, C). This reanalysis notably identified many previously undetected genes, including highly significant examples, with clear repeating oscillations and high relative amplitude that the prior work (Figure 3B, D). The genes in the respective light and temperature-entrained 2-day datasets were largely unique (consistent with the prior work), with only 96 being identified in both, fewer than would be expected by chance (Fisher’s Exact *p*-value =1, data not shown).

**Figure 3:**
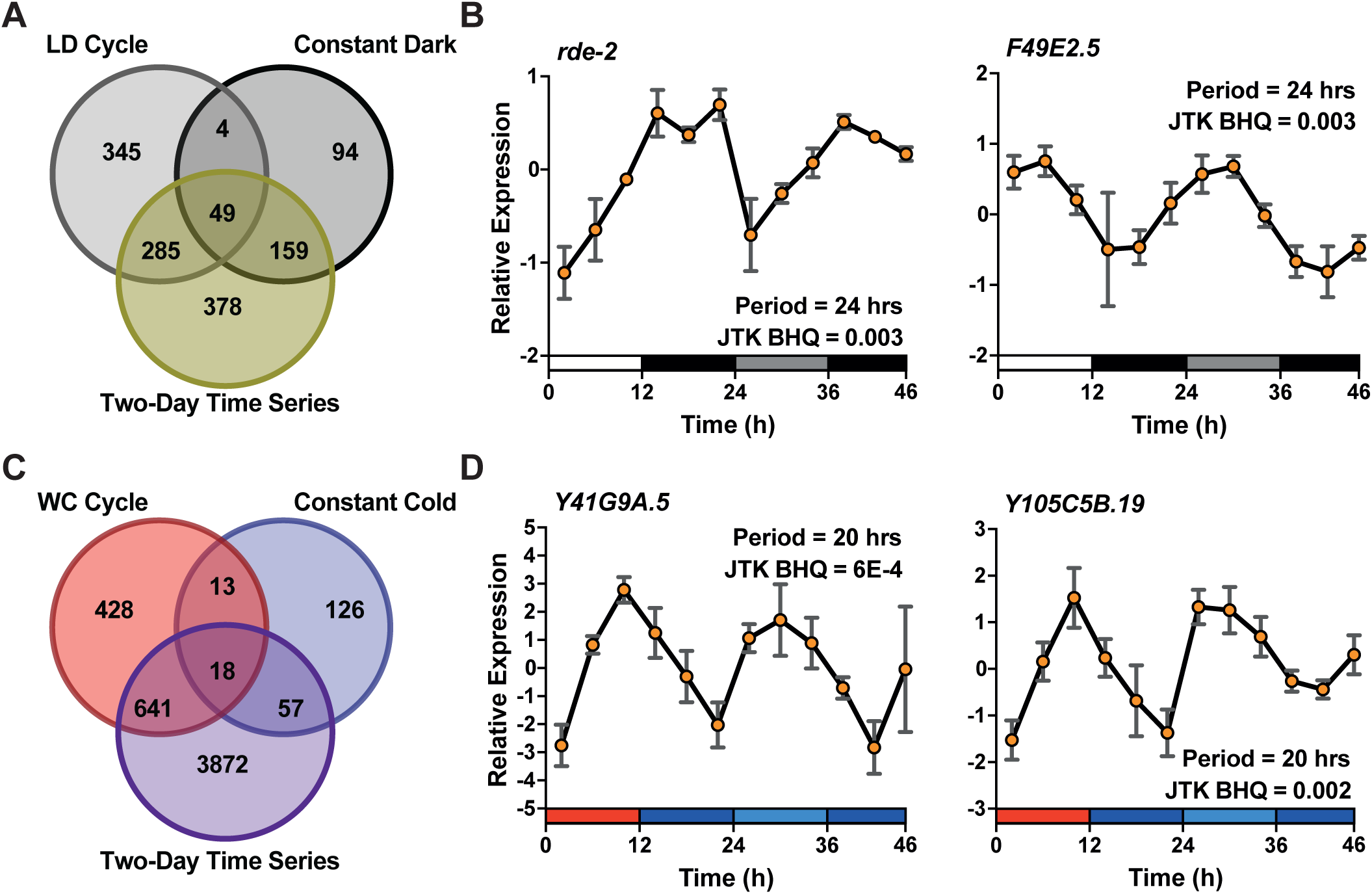
JTK_Cycle reanalysis of the 2010 Dataset. **A** and **C:** Venn diagrams showing the number of genes collectively and individually identified by JTK_Cycle (BHQ < 0.05) under a 12:12-hour environmental cycle, constant conditions and over full two-day time series following light (**A**) and temperature (**C**) entrainment protocols. **B** and **D:** Examples of significantly rhythmic genes identified by JTK_Cycle (both within the lowest 20 BHQ values) over the full two-day light (**B**) and temperature-entrainment (**D**) time series. Figures include period estimates and BHQ values as generated by JTK_Cycle. Shown are average expression values of 3 (**C**) or 3-5 (**D**) replicates ± SEM. Relative expression values were calculated previously as described [14]. Time 0 represents the time of light onset or temperature increases respectively with environmental conditions indicated by boxes along the *x*-axis (**B:** white = light, black = dark, grey = subjective day (dark), all at 18°C. **D:** red = 25°C, blue = 15°C, light blue = subjective day (15°C)).

### Convergence in period and phase between transcriptomic datasets

As evident from the analyses above and work in more established chronobiology models, circadian analysis methods for time-series data can result in inconsistencies in the number and identities of rhythmic genes, which can vary by dataset, analysis method and significance thresholds [29,78,85–87]. Published datasets do often converge however, on TTFL-associated or select other clock-controlled genes, including with respect to the phase of oscillation [7,88]. As such, we compared datasets with a view that overlapping genes, or those that differ could be informative in moving towards a mechanistic understanding of circadian biology in *C. elegans*.

The two-day analyses of the 2010 time series (ANOVA *p* < 0.05 and JTK_Cycle, BHQ < 0.05) showed limited coherence in gene identities with the RNA-seq time series: 28 light-entrained and 75 temperature-entrained genes were shared with the RNA-seq dataset (Figure 4A, D; Table S7). However, we found this coherence to be statistically significant for both light and temperature comparisons (Fisher’s exact test *p*-value < 0.05), reflecting a modest increase over expected convergence by random chance (17 and 51 genes respectively). A total of 8 of these genes were identified in all three datasets: *T06A1.5*, *C06H5.6*, *C37C3.2*, *H06I04.6*, *smd-1*, *vha-5*, *xbp-1* and *Y54G11A.7*. To further compare these datasets, we noted that of the 95 overlapping light or temperature-entrained genes, approximately 40% were assigned the same period by JTK_Cycle (Figure 4B, E; JTK_Cycle was allowed to assign periods of 20, 24 or 28 hours). Of these genes, most were similar in JTK_Cycle-assigned phase, and showed comparable expression patterns (examples given in Figures 4B, E) [7,88]. As such, these numerically modest subsets of genes (highlighted in Table S7) likely represent the strongest candidates from our work for either mechanistic involvement in circadian timing in *C. elegans* or for use as reporters of the circadian system.

**Figure 4:**
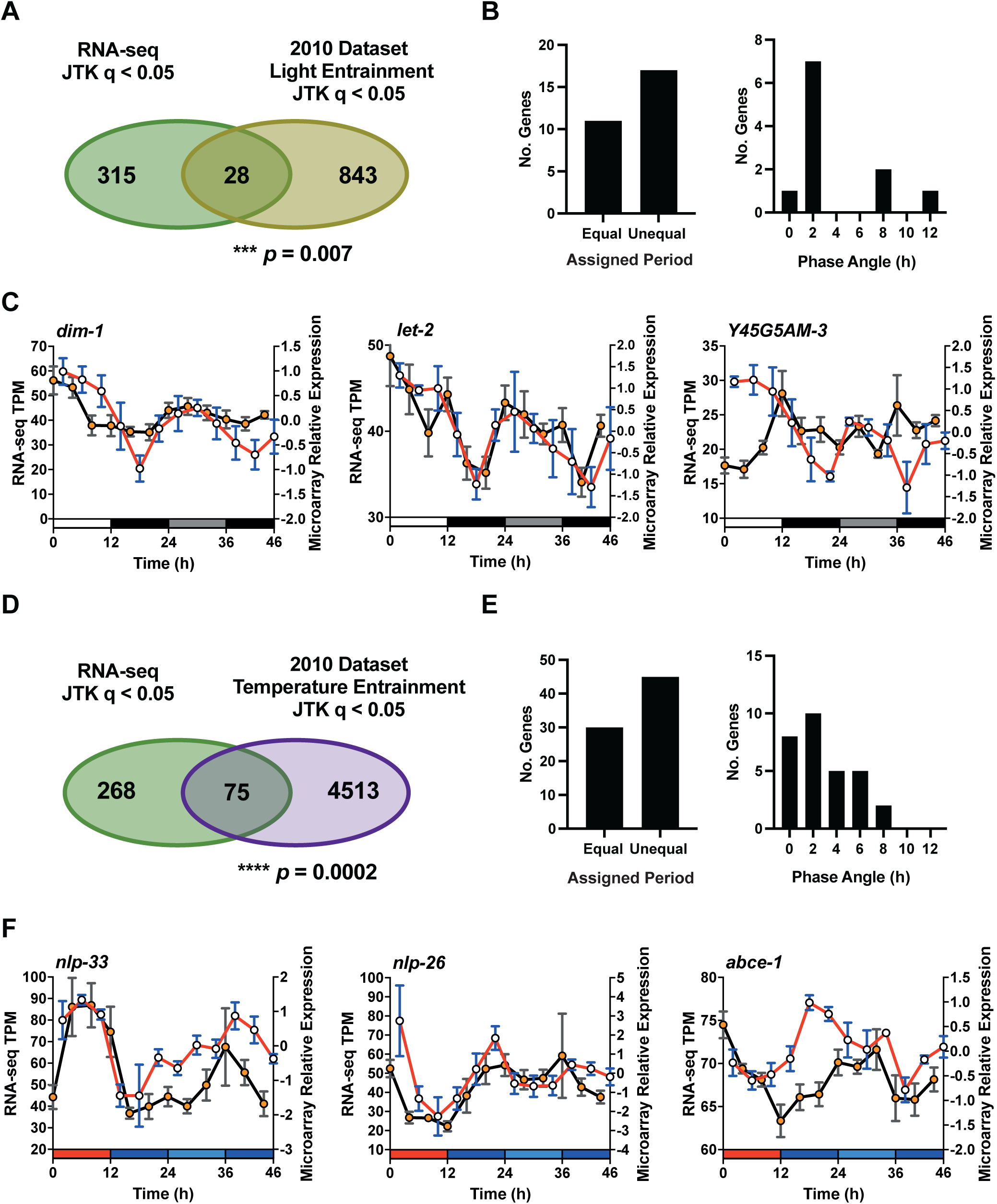
Comparisons of gene identities, periods and phases between the RNA-seq and 2010 datasets. **A** and **D:** Venn diagrams showing the numbers of genes collectively identified by JTK_Cycle (BHQ < 0.05) under over our full two-day RNA-seq time series and the 2010 microarray dataset under light (**A**) and temperature (**D**) entrainment protocols. *p*-values calculated from respective Fisher’s exact tests comparing significantly rhythmic against all remaining genes detected in both datasets. **B** and **E:** Numbers of genes from **A** and **D** respectively with equal and unequal JTK_Cycle-assigned periods. JTK_Cycle-assigned phases are shown for the proportion of genes with equal periods. **C** and **F:** Examples of expression patterns from overlapping genes in **A** and **D** respectively, giving two phase-coherent and one phase incoherent example in each case. Shown are average expression values of replicates ± SEM. For the 2010 dataset, relative expression was calculated previously as described [14]. Time 0 represents the time of light onset and/or temperature increases respectively in both datasets with environmental conditions (here specific to the 2010 datasets) indicated by boxes along the *x*-axis (**C:** white = light, black = dark, grey = subjective day (dark), all at 18°C. **F:** red = 25°C, blue = 15°C, light blue = subjective day (15°C)).

Further to the above, considering specific genes of interest we generally saw a lack of rhythmicity in TTFL related homologues in the 2010 dataset (consistent with the published analysis and our RNA-seq data) and other specific genes of interest (Figure S1, full 2010 JTK_Cycle results in Table S6). The *per* homologue, *lin-42* did not show a clear response to environmental cycles across all three time series, nor did *nhr-23*, homologous to mammalian *Rora*/*Rorb* genes and reportedly required for circadian rhythms in *C. elegans* [29]. One exception is *atf-2* which was rhythmic in a light:dark cycle only. The *pdfr-1* gene, which was highly autocorrelated in our data, showed no consistent expression between datasets. We also did not see concordant rhythmicity two previously described *C. elegans* circadian reporter genes: *sur-5* [66,89], which showed no rhythm in any dataset, nor *nlp-36* which was rhythmic in the 2010 study [14], and remained so in the reanalysed temperature-entrained data only. A final gene of interest, *xbp-1*, whose mammalian homologue has been associated with 12-hour ultradian expression rhythms [90–93], was rhythmic across all three datasets. This gene did show consistent expression in temperature-entrained and RNA-seq dataset, particularly during the entrainment phases (Figure S1).

### WGCNA reveals genes co-expressed coherently with environmental cycles

To further analyse our RNA-seq data, we sought to investigate prevailing gene expression trends without prior assumptions as to the expression patterns (such as the sinusoidal expression detected by JTK_Cycle or the assumed 24-hour periodicity in autocorrelation). To do this we employed weighted gene co-expression network analysis (WGCNA) [75], a pairwise correlation and hierarchical clustering-based approach that has previously been used to identify varied circadian expression patterns in mouse and human data [94,95]. Taking our full dataset of 16,176 expressed genes (> 0 TPM in > 50% of samples; Figure 1), performing WGCNA generated 13 modules (clusters) of similarly or co-expressed genes (shown in Figures 5A, 6A and S2; full list of assignments in Table S8). Each module is represented by an eigengene expression pattern, which represents the values of the first principal component in each sample, here averaged at each timepoint. These modules should thus indicate the most prevalent underlying expression trend of the genes within.

**Figure 5:**
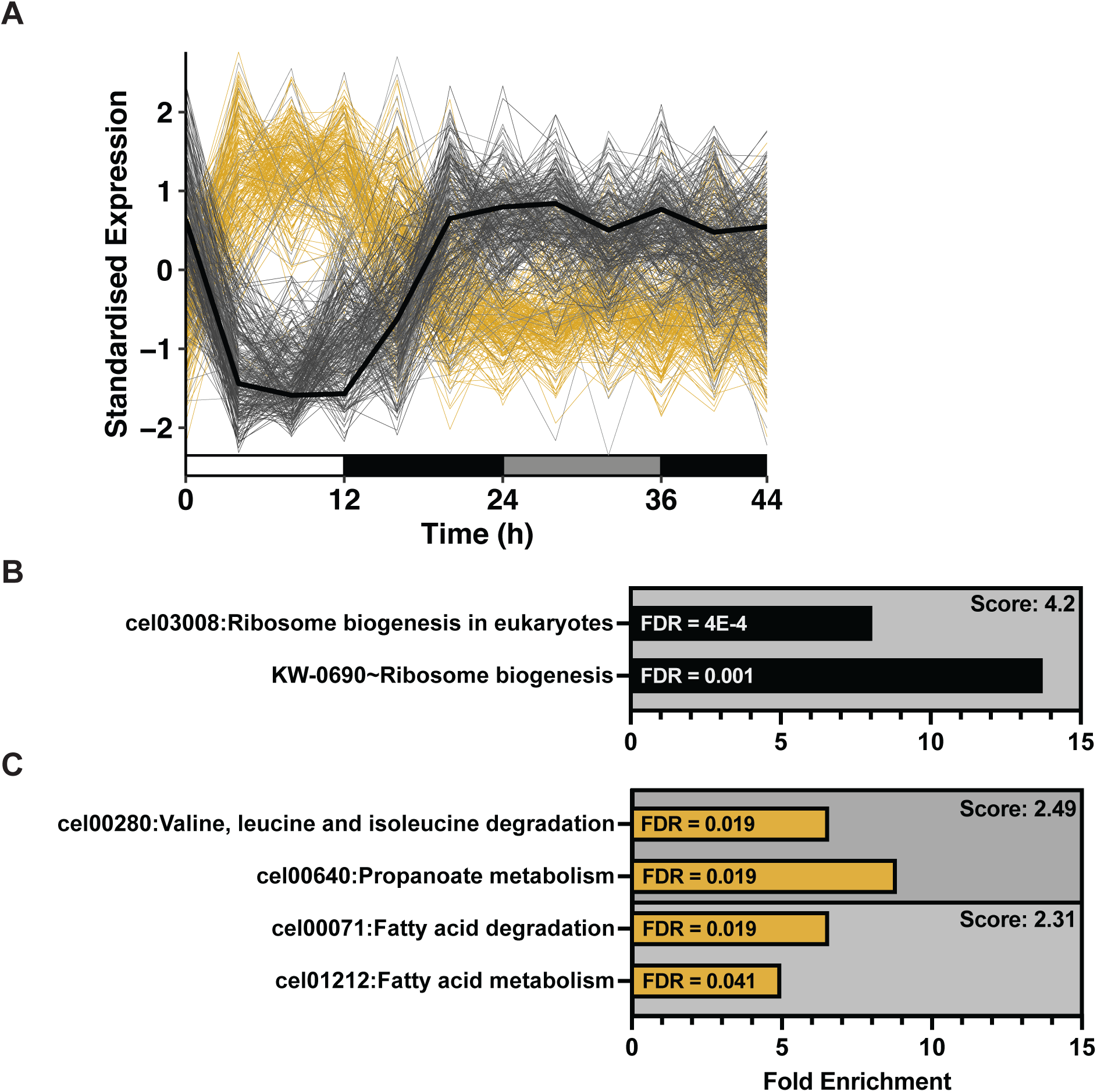
WGCNA analysis reveals genes co-expressed in-phase with a light and temperature cycle. **A:** Standardised expression of all 511 genes in the Black module over time. Black line represents the module eigengene values, averaged by timepoint. Grey lines indicate average standardised expression of the 300 genes with the least overall deviation from the eigengene and gold lines represent the remaining 211. Darker lines indicate closeness to the eigengene in groups of 50. Bars along *x*-axis indicate environmental conditions: white = light/20°C, dark grey = dark/15°C, light grey = subjective day (dark/15°C)). **B:** DAVID enrichment analysis of genes in Black module subsets. Here we are showing fold enrichment and FDR values for significantly enriched terms, which DAVID has groups into enrichment clusters. Each cluster has an associated enrichment score calculated from the geometric mean of the *p*-values of the terms within (in -log scale such that higher enrichment scores represent greater average *p*-value significance). For simplicity, we are here showing clusters for which > 50% of terms within had FDR < 0.05). Full list in Tables S9 and S10.

Of particular interest to our study, one WGCNA module, designated ‘Black’, revealed gene expression changes coherent with our environmental light and temperature cycle. This module contained 511 genes, represented by an eigengene that declines through light and warm conditions, and increases at the onset of the dark and cold phase (Figure 5A) while showing much reduced variation in constant conditions. This module also identified many genes following the opposite trend increasing from a lower baseline in response to light and increased temperature), indicating two antiphasic expression trends corresponding to (and likely driven by) the environmental cycle (Figure 5A). Given that both of these trends represented real gene expression patterns, we divided the module and investigated these trends separately. To do this, the standardised expression of each of these 511 genes was ranked by summed deviations from the eigengene over the time series. We then generated arbitrary subsets of the 300 least-deviating genes and the remaining 211 (plotted in grey and gold respectively in Figure 5A). In doing so we highlight genes identified by WGCNA that cycle daily and with circadian periodicity, in response to cycles of light, temperature or both.

To investigate the prevailing functions of these light and temperature-cycling genes, we performed functional annotation analysis on the two Black module subsets independently. This highlighted ribosome biogenesis as a metabolic process associated with genes downregulated in our light and warm conditions (grey genes in Figure 5A; Figure 5B; Table S9), and functions including fatty acid and amino acid metabolism as potentially upregulated in light and warm conditions (gold genes in Figure 5A; Figure 5B; Table S10), in both cases reidentifying similar annotations to those found for the JTK_Cycle analysis (Figure 2C). One TTFL-associated gene was assigned to this module, the previously discussed *pdfr-1* (Table S3, Figure S1).

### WGCNA reveals high numbers of co-expressed genes with 16-hour periodicity

Distinct from our circadian analyses, the most striking, and novel, expression pattern identified by WGCNA were genes co-expressed with 16-hour ultradian periodicity. These patterns occur over multiple cycles and are exemplified by the Brown and Yellow modules (Figures 6A and S2). The Brown module contained 1244 genes, including many with expression patterns closely represented by that of the eigengene (300 of which are highlighted in Figure 6A, exemplifying that the module eigengene reflects the expression patterns of the genes therein). As well as standardised expression, TPM values similarly conformed to the 16-hour eigengene trend (with the 12 genes least-deviating from the eigengene plotted in Figure 6B). This included highly abundant genes with > 2-fold changes in expression. Given the difference in period with our environmental cycles, the relationship of these expression patterns with our entrainment protocol is unclear, but the initial peak of these genes and the module eigengene notably occurs at time 0 (the beginning of the light/warm phase).

**Figure 6:**
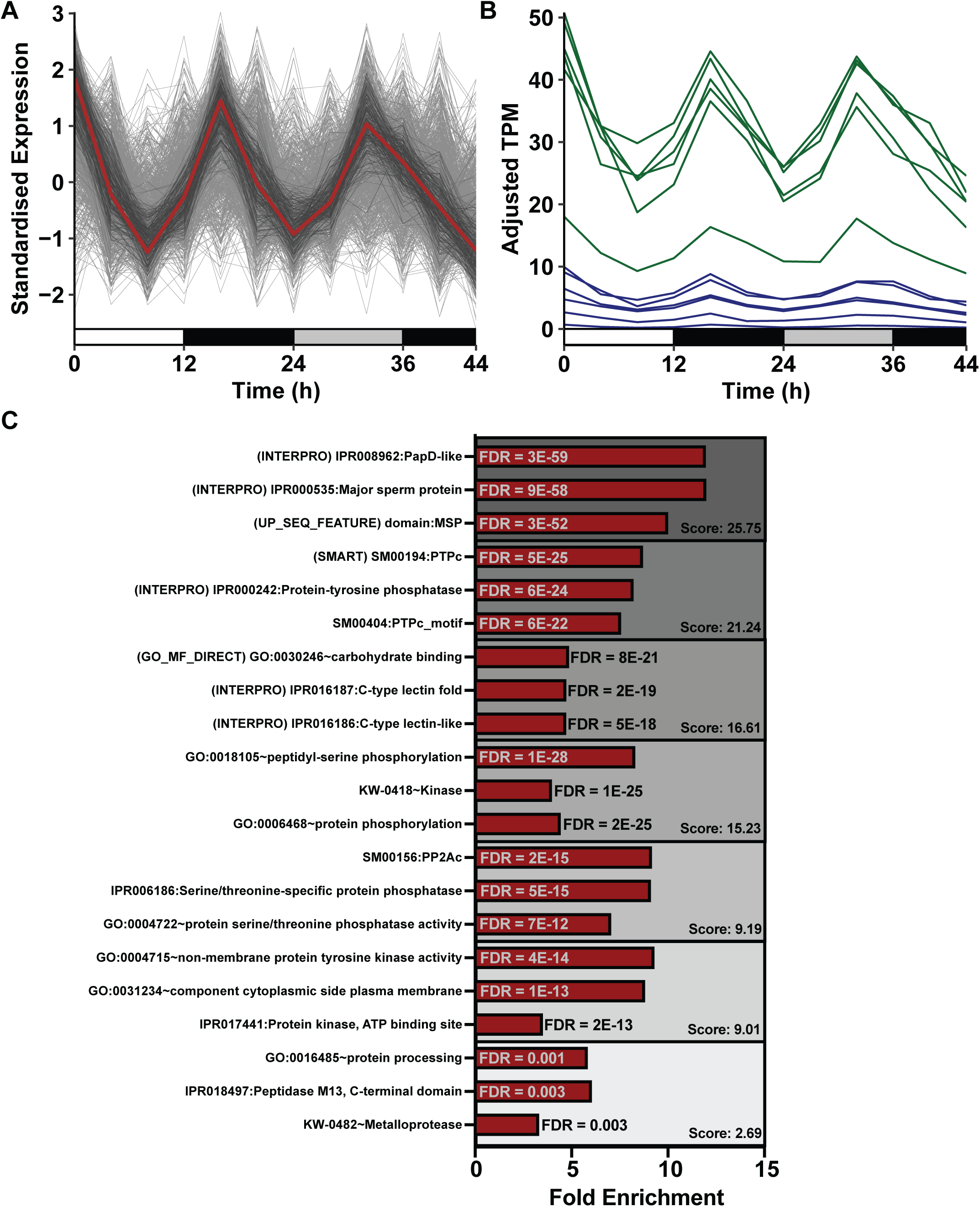
WGCNA clustering reveals genes with 16-hour periodicities associated with multiple functions. **A:** Standardised expression of all 1244 genes in the Brown module over time. Brown line represents the module eigengene values, averaged by timepoint. Grey lines indicate average standardised expression of all 1244 genes, with darker lines indicating the 300 genes with the least overall deviation from the eigengene in groups of 50. **B:** The 12 genes with the smallest total deviation from the Brown module eigengene in terms of standardised expression, here expressed in Batch-adjusted TPM (see Methods), with each replicate averaged by timepoint. *msp* genes are shown in green, with other genes in blue. Bars along *x*-axis indicate environmental conditions: white = light/20°C, dark grey = dark/15°C, light grey = subjective day (dark/15°C)). **C:** DAVID functional annotation of genes in Brown module. Shown are DAVID enrichment clusters and Fold enrichment and FDR values for the three most significant terms within, and the associated enrichment score for each cluster (based on the inverse of the geometric *p*-value mean). Enrichment clusters are shown here if FDR < 0.05 in > 50% of terms within the cluster.

DAVID analysis of the Brown module suggested an array of functions associated with the genes within, with far more significantly enriched terms being identified than in our JTK_Cycle and Black module analyses (Figure 6C, Table S12). This is reflected in much greater enrichment scores of the clusters shown in Figure 6C (which scale inversely to the average *p*-values of the terms within). The most enriched cluster included genes of the major sperm protein (MSP) gene class. Genes encoding an MSP domain were also among the genes most closely represented by the module eigengene trend; six of the 12 traces shown in Figure 6B represent transcripts mapped to *msp* genes (plotted in green). Conversely, many other highly enriched functions not associated with MSP were identified; terms contained within the six remaining enrichment clusters shown in Figure 6C (with FDR < 0.05 in > 50% of terms) including a number of kinase and phosphatase-related terms (Figure 6C). These two enrichment clusters in particular notably contained very few (< 3) *msp* genes. This analysis therefore suggests hitherto unreported 16-hour oscillatory co-expression of a large subset of *C. elegans* genes, associated with broad and essential cellular and organismal functions.

## Discussion

In this work, we investigated oscillatory gene expression in *C. elegans* hermaphrodites post-development, performing RNA-seq on samples collected over a circadian time series. Using a widely applied method in JTK_Cycle [6,7,76–79] coupled with autocorrelation analysis, we identified evidence for daily oscillations in gene expression in many previously unreported genes [14,29]. However, we also noted that our ability to detect rhythms was considerably poorer in the absence of environmental stimuli. A comparable reanalysis of previously published time series data (‘the 2010 dataset’; [14]) was consistent with this observation, while suggesting the persistence of considerably more rhythmically expressed genes, particularly following cycles of temperature. Further analysis of our RNA-seq dataset using WGCNA revealed prevalent co-expression patterns coherent in phase with our environmental cycles and strikingly, thousands of genes with high amplitude 16-hour co-expression patterns.

### Daily rhythms in *C. elegans* identified by RNA-seq

With respect to circadian gene expression rhythms in *C. elegans* and their potential functions, we identified hundreds of novel oscillating genes over a two-day time series (543 with JTK_Cycle BHQ < 0.05 or autocorrelation r > 0.7; Figure 2, Table S1). Functional annotation of the JTK_Cycle results suggested fatty acid metabolism and ribosome biogenesis as prevailing physiological outputs encoded by these genes, both of which could represent rhythmic processes conserved with mammals [96,97]. Ribosome biogenesis could represent a particularly essential conserved function, having been reported to be circadian clock-regulated in-part at the mRNA level in mammals [98]. The turnover (synthesis and degradation) of ribosomes, representing 3-6% of total cellular protein [99–101], has also been shown to exhibit a circadian rhythm in mammalian cells [102]. Several of our rhythmic ribosome biogenesis-associated genes in *C. elegans* are involved in the processing of ribosomal RNAs [103], orthologues of which exhibit robust mRNA circadian oscillations in other species, including orthologues of *nol-58*, *Y66H1A.4* (*garr-1*) and *xrn-2* in mammalian livers [104,105] and *nol-58* in the circadian model fungus, *Neurospora crassa* [78,106]. Rhythms in rRNA itself would by design, not be detected in our mRNA sequencing, but if shared with other models would reflect conservation of daily timing in a critical, energetically expensive aspect of cellular homeostasis.

While potentially physiologically relevant, the enriched functions we detect may reflect environmentally driven oscillations more so than endogenous timekeeping; we notably detected considerably more significant cycling genes using JTK_Cycle over an environmental cycle alone, but very few in constant conditions (Figure 2D; Supplementary Tables 1, 4, 5). Our WGCNA analysis also identified hundreds of co-expressed genes with clear 24-hour expression patterns under a light and temperature cycle, while varying much less in constant conditions (the Black module; Figure 5). Interestingly, this module also showed significant enrichment for fatty acid metabolism and ribosome biogenesis-related ontology terms. These respective sets of genes were also in antiphase, so likely did not simply reflect the varying rates of metabolism in our poikilothermic model. That we identified these terms using differing complementary approaches strongly suggests the validity of daily cycling of genes involved in these processes. However, it does highlight the distinction between rhythms under environmental cycles, as reported in some circadian transcriptomics literature [87,107], and endogenous timekeeping, leaving open the question of whether *C. elegans* is capable of sustaining endogenous rhythms.

### Comparative analyses and the *C. elegans* circadian system

Our reanalysis of the 2010 dataset [14] revealed a similar reduction in cycling genes in constant conditions relative to environmental cycle data, but identified far a greater proportion of genes to oscillate overall (∼25%; Figure 3, Table S6). It is unclear why we saw such stark differences between datasets, with the respective experiments being similar with respect to sampling method, temporal resolution and *C. elegans* strain (N2). One potentially pertinent difference could be entrainment differences, with the 2010 work applying light and temperature cycles independently and over a wider temperature range. A previous RT-qPCR reanalysis also could not recapitulate rhythms in select 2010 dataset genes following cycles of colder temperatures [15]. The 2010 study itself reported rhythms to co-occur in only two genes in the respective light and temperature datasets [14] (rising to 96 in our reanalysis, ∼1.7% of detected rhythmic genes and fewer than expected by chance. *C. elegans* has also been reported to optimally entrain its gene expression and locomotor activity to light and temperature cycles with light coinciding with lower temperatures [16,22,66], contrasting with our approach. Importantly however, while *C. elegans* entrainment remains an important question, variability in the number and identities of rhythmic genes is also a common observation (and fundamental concern) in circadian transcriptomics literature. This includes variation by analysis approach, or when applying the same analysis to different datasets [7,29,78,85–88]. Ultimately our work emphasises the need for cautious experimental methodology and analysis choices when studying relatively non-robust processes like *C. elegans* circadian timing.

Consistent between RNA-seq and 2010 datasets, we detected limited evidence of rhythmicity in *lin-42* or other TTFL-related *C. elegans* sequence homologues [27,28] (Tables S3, S6), as in prior literature [14,15,29]. We also saw no clear or comparable response to environmental cycles on *lin-42* expression across datasets (Figure S1), nor was it assigned to a 16-hour WGCNA module (Table S3). This result, repeatedly observed in *C. elegans*, contrasts with conserved TTFL models of transcriptional circadian rhythms, which are defined by oscillating mRNA and protein abundance of evolutionarily conserved *per* homologues [1,12]. While *lin-42* deficiency has been shown to affect circadian period [66], it is most well studied as a heterochronic gene, important for normally timed moulting within *C. elegans*’ temporally defined developmental program [41–47], along with several other TTFL homologues [29,42,47]. *lin-42* expression is much greater during development than in adults [40], where it does oscillate along with 2000-4000 other genes in *C. elegans* and other nematodes coincident with moulting [33–36,38–40]. As such, *lin-42*’s prevailing role appears to be one that parallels, but is distinct from, circadian timing.

The mechanisms underpinning transcriptional rhythms are beyond the scope of our work, but convergence between datasets analysed here could yield insights into robust or important oscillating genes. Of the 95 genes identified by JTK_Cycle across both RNA-seq and 2010 datasets, a high proportion were similar in phase (Figure 4, Table S7). Given that circadian transcriptomics studies typically converge on TTFL genes of similar phases in other models [7,88], this subset could contain candidates of mechanistic importance.

Conversely, posttranscriptional circadian rhythms can be observed in the absence of any transcription in some systems [108,109]. In *C. elegans* itself, abundance rhythms in the protein GRK-2 have been reported to persist without a corresponding mRNA oscillation [15]. Further, gene and protein abundance rhythms have been found to be present, if sometimes less robust, when the TTFLs are perturbed (including in *Drosophila* lacking *per* [6]) and mammalian systems lacking *Bmal1* [110,111] or *Cry1*/*Cry2* [112,113]). In this context, and given that we did not detect rhythms in previously described reporter genes (*nlp-36* and *sur-5*), our reproducibly rhythmic genes could have utility as reliable transcriptional outputs, for understanding circadian rhythms when a conserved transcriptional oscillator is potentially absent.

### Novel 16-hour periodic co-expression in *C. elegans*

The most abundant expression patterns emerging from our RNA-seq analyses were not circadian rhythms, but high-amplitude, 16-hour co-expression of genes identified by WGCNA [75] and represented by the Brown and Yellow modules (Figures 6, S2). Over 1000 genes (∼7.5% of detected genes) were assigned to the Brown module, with many being well-represented by the eigengene expression profile. Ultradian (< 24-hour) rhythms have been widely reported in nature, including 12-hour harmonics of the circadian rhythm in mouse and human transcriptomic data [92,104] and also in *C. elegans* (again from the 2010 dataset) [93]. Circatidal rhythms, rhythms of a similar periodicity synchronised by tides, can also be seen in marine organisms across phyla, in some cases being contingent on expression of TTFL-essential genes and proteins [114–119]. However, the 16-hour co-expression we observe is distinct in period from these harmonics along with other shorter periodic events in *C. elegans* including defecation [120,121], egg laying [122] and pulsatile developmental gene expression [123]. We also saw limited assignment of TTFL homologues to these modules (although the TTFL-associated *ces-2* and *atf-2* were notably assigned to the Yellow module (Table S2)).

One function of this 16-hour co-expression trend likely pertains to reproduction. Overwhelmingly, the most enriched terms identified by functional annotation related to genes of the major sperm protein (MSP) class (Figure 6C; Table S11). MSP genes were also among those most closely aligned with the module eigengene expression profile (Figure 6A, B). MSPs are nematode-specific proteins expressed highly and exclusively in sperm [124]. They perform dual roles, as key cytoskeletal components of spermatozoa, providing the basis for motility [125], and as secreted proteins necessary for the meiotic maturation of oocytes [126]. Given our experiment used FUDR, a DNA synthesis inhibitor, to prevent reproduction, meiosis itself could not have been taking place in our experiment. These 16-hour co-expression profiles may therefore reflect an underlying transcriptional mechanism by which oocyte maturation, ovulation or fertilisation are temporally coordinated [127].

Importantly however, many of our Brown module genes did not have MSP or direct reproduction-related annotations; DAVID enrichment analysis of the Brown module resulted in higher enrichment scores and far more significant enrichment terms than JTK_Cycle or Black module gene lists (Figure 6C, Table S11). Multiple enrichment clusters relating to protein phosphorylation were particular cellular functions associated with Brown module 16-hour oscillatory genes. These terms could therefore reflect much wider temporal regulation of *C. elegans* biology post-development.

Collectively, our transcriptomics analyses provide insights into temporal coordination in *C. elegans*, highlighting both the peculiarities of its circadian system, but also offering a lens through which we can appreciate the broader landscape of rhythmic phenomena across diverse organisms. *C. elegans*’ primary utility as a chronobiology model may be, for example, in understanding how rhythms are generated in the absence of a conserved transcriptional oscillator. The unique 16-hour co-expression we observe, predominantly associated with msp genes, hints at a potential link between well-studied reproductive timing and temporal coordination in post-developmental nematodes. Finally, this work emphasises the value of considered data collection and analysis when considering non-robust or noisy biological processes like transcriptional oscillations; we highlight the disparities that can emerge using established algorithms like JTK_Cycle as well as firmly establishing WGCNA as an effective tool for identifying large scale co-expression in time series - omics data.

## Methods

### Nematode maintenance

All work was performed using N2 strain nematodes, reared on plates seeded with OP50 strain *Escherichia coli*, both of which were provided by the Caenorhabditis Genetics Center (CGC; https://cgc.umn.edu), which is funded by NIH Office of Research Infrastructure Programs (P40 OD010440). Where applicable, plates containing 25 μM 5-Fluorodeoxyuridine (FUDR), were twice seeded by adding 10X culture and UV-killing *E. coli*.

### RNA-seq entrainment protocol and RNA harvesting

Entrainment conditions consisted of 12:12-hour cycles of light (10 μmol m^−2^ ^s−1^) at 20°C and darkness at 15°C. Parental generation larvae were reared under entrainment conditions throughout development. Age-synchronised progeny were obtained by bleaching parental hermaphrodites [128] and maintaining eggs/L1 larvae on unseeded plates for 2 days. Nematodes were transferred to seeded plates to develop for 2 days and then transferred again to plates containing 25 μM 5-Fluorodeoxyuridine (FUDR) to prevent reproduction and interference from developing larval gene expression [129]. To transfer of nematodes to new plates in experiments, plates were washed by adding 2-3 mL of sterile S buffer or H_2_O. All manipulations were performed around the start of the light/20°C warm phase (dawn). Nematodes were entrained for a further 3 days and then constant darkness at 15°C for 1 day. Entrainment and subsequent constant conditions were accomplished in a single growth chamber, in which warming took approximately 85 mins ± 10 mins and cooling took 55 minutes ± 5 mins.

RNA Collections took place starting at dawn (time 0) and then followed every subsequent 4 hours for 2 days (12 timepoints in total). Each biological sample comprised approximately 150 hermaphrodites housed on an individual NGM plate. Nematodes were harvested by washing in 2 mL S buffer, centrifugation for 1 min, aspiration to approximately 100 μL, resuspension in 250 μL TRIzol Reagent (Ambion), mixed by pipetting, and immediately frozen at -70°C. Three samples were collected simultaneously at each timepoint and 2 independent time series utilising the same conditions were performed, resulting in 6 replicates in total.

### RNA processing and sequencing

Samples for RNA-seq were processed by batch in randomised order in sets of 12. To extract RNA, samples in TRIzol were defrosted then subjected to 3 liquid nitrogen freeze/thaw cycles. Subsequently, samples were left to sit for 5 minutes at room temperature, 50 μL chloroform was added and samples were centrifuged for 15 minutes at 13200 RPM at 4°C. Most of the aqueous phase was transferred to a new tube and then processed following the protocol of the QIAGEN RNeasy Micro kit, including on-column RNase-Free DNase treatment.

Sample quality was checked using the Agilent 2100 Bioanalyzer on RNA Nano Chips, with a majority having a RIN value of 10. 100 ng total RNA for each sample for library preparation using the NEBNext RNA Ultra II RNA directional library prep kit. Unique 8 bp dual indices were added to each sample. Sample quality was again checked using 2100 Bioanalyzer. Sequencing was then performed across 2 lanes on an Illumina HiSeq 3000 machine.

### RNA sequence processing

Sequence quality was checked by MultiQC software [130]. Sample depth ranged from 12.4 to 32.8 million reads per sample, except for one sample at the 44-hour timepoint, which contained very few reads and was excluded from further analysis, resulting in 5 replicates at this timepoint (71 samples in total). Adapter sequences were trimmed using Cutadapt [131], and samples reanalysed using MultiQC. Salmon [132] was used for quasi-mapping of sequencing reads to the *C. elegans* genome (assembly WBcel235, obtained from GenBank [133], and for quantification of gene expression. TPM values were obtained using Sleuth [134], within R Studio [135,136], following preparation of files using wasabi (COMBINE-lab, 2018).

### RNA-seq analysis: JTK_Cycle, autocorrelation, WGCNA and DAVID functional annotation

Data was initially filtered to include only genes with > 0 TPM in ≥ 36 of all 71 samples. These full tie series data are given in Table S12 One-way ANOVA was performed on each detected gene in R to pre-screen the data for significant changes in expression (*p* < 0.05). This was performed independently for both the whole filtered dataset and dividing the dataset into 2 sets of 6 timepoints representing the entrainment day and subsequent day.

Rhythmic genes were detected using the JTK_Cycle function in the MetaCycle R package [74], setting period limits to 20 and 28 hours for the full time series and 20 to 24 hours for individual days. Genes with a Benjamini-Hochberg *q*-value < 0.05 were considered significantly rhythmic. For period and phase comparisons, differences were calculated using JTK_Cycle-assigned values. We also applied ANOVA to log_2_(TPM+1) values and a non-parametric Kruskal Wallace as alternate pre-screening approaches prior to JTK_Cycle analysis on to the two-day time series data, and found that this made only minor differences in resulting rhythmic gene lists (in the case of the former, reidentifying all but 3 of 343 genes with BHQ < 0.05, and expanding the list by 43, none of which were highly significant).

Autocorrelation analysis was performed in Microsoft Excel, ranking expression over the entrainment and constant conditions days and calculating Spearman’s rank coefficient (r_s_) between timepoints spaced 24 hours apart.

WGCNA analysis was performed using the WGCNA R package [75] to analyse the filtered dataset (> 0 TPM in > 50% of samples). Samples were first corrected for batch effects using ComBat within the sva R package [137,138]. WGCNA was applied generally following settings and methods used in the online tutorial (section I); first clustering by sample (resulting in 11 of 71 being excluded as outliers). An adjacency matrix of all genes across the remaining samples was then generated based on their pairwise co-expression (absolute Pearson correlation coefficient). The soft-threshold power was set manually at 6 such to minimise the numbers of genes assigned to no module (Grey) and the largest module (Turquoise). To investigate gene expression over time, eigengene values were averaged by timepoint and compared to standardised expression of the genes within the module (subtracting a mean and dividing by standard deviation).

Functional annotation Clustering was performed using the Functional Annotation Clustering from Database for Annotation, Visualization and Integrated Discovery (DAVID, version: DAVID 2021) [80,81], including all annotation categories. For all analyses we compared our gene list of interest to a background list of our initial 16,716 genes with detectable expression (TPM > 0 in > 50% of samples). Some terms were edited for ease of presentation in figures. Where presented, the enrichment score represents the geometric mean (-log scale) of the *p*-values within a cluster calculated by DAVID.

### Reanalysis of 2010 dataset microarray data

Processed, normalised and standardised data (as described [14]) were obtained from Gene Expression Omnibus (GEO; www.ncbi.nlm.nih.gov/geo/; accession: GSE23528). To compare this data with ours, Affymetrix IDs were converted to Official Gene Symbol using the DAVID Gene ID conversion tool [80,81]. Official Gene Symbols were retrieved for 21318 of 22625 initial transcripts.

Light and temperature time series data were separated and analysed as independent time series, including three replicates for all conditions except the warm:cold cycle which included five. All transcripts were analysed by ANOVA and JTK_Cycle as a two-day time series and individual six-point series as above, with duplicate and unnamed transcripts being removed from counts of rhythmic genes. Venn diagram comparisons were analysed using one-tailed Fisher’s exact tests, comparing genes with ANOVA *p* < 0.05 and JTK_Cycle BHQ < 0.05 against all unique detected genes common to both datasets (TPM > 0 in 50% of samples RNA-seq data, all retrieved genes in microarray data; 13,656 common genes). Expected values were calculated from contingency tables. Phase angles were calculated from differences in JTK_Cycle-assigned phases, which represents the timepoint at which the acrophase of the fitted curve occurs. We adjusted for circular phase using the intrinsic JTK_Cycle-assigned period in each case (20, 24 or 28hrs).

### Data analysis and figures

Data were analysed using Microsoft Excel Microsoft Excel (Microsoft Corporation), Prism GraphPad 9/10 (Boston Massachusetts USA) and R Studio [135,136]. All figures were plotted using Graphpad Prism, Adobe Illustrator (Adobe Inc.) or ggplot2 in R Studio [139].

## Acknowledgements

We thank Amanda Davis for helpful discussions on study design and execution, John Davey for bioinformatics support, John O’Neill for analysis advice and members of the Chawla and Davis Labs for helpful discussion. This work was funded by a Departmental PhD studentship to JM, BBSRC Grant (BB/N018540/1) to SJD and pump-priming funds from the Department of Biology, University of York.

## Author contributions

JM, SJD and SC designed research; JM, LG, SRJ performed research; JM, KN analysed data; JM wrote the paper with editing contributions from all authors.

The authors declare no competing interests.

## References

1. Patke A, Young MW, Axelrod S. 2020 Molecular mechanisms and physiological importance of circadian rhythms. Nat Rev Mol Cell Biol 21, 67–84. (doi:10.1038/s41580-019-0179-2)

2. Dunlap JC, Loros JJ. 2017 Making Time: Conservation of Biological Clocks from Fungi to Animals. Microbiol Spectr 5, 1–32. (doi:10.1128/microbiolspec.funk-0039-2016)

3. Harmer SL. 2009 The circadian system in higher plants. Annu Rev Plant Biol 60, 357–377. (doi:10.1146/annurev.arplant.043008.092054)

4. Liu Y, Tsinoremas NF, Johnson CH, Lebedeva N V., Golden SS, Ishiura M, Kondo T. 1995 Circadian orchestration of gene expression in cyanobacteria. Genes Dev 9, 1469–1478. (doi:10.1101/gad.9.12.1469)

5. Eelderink-Chen Z, Bosman J, Sartor F, Dodd AN, Kovács ÁT, Merrow M. 2021 A circadian clock in a nonphotosynthetic prokaryote. Sci Adv 7, 1–7. (doi:10.1126/sciadv.abe2086)

6. Hughes ME, Grant GR, Paquin C, Qian J, Nitabach MN. 2012 Deep sequencing the circadian and diurnal transcriptome of Drosophila brain. Genome Res 22, 1266–1281. (doi:10.1101/gr.128876.111)

7. Zhang R, Lahens NF, Ballance HI, Hughes ME, Hogenesch JB. 2014 A circadian gene expression atlas in mammals: Implications for biology and medicine. Proc Natl Acad Sci U S A 111, 16219–16224. (doi:10.1073/pnas.1408886111)

8. Konopka RJ, Benzer S. 1971 Clock mutants of Drosophila melanogaster. Proc Natl Acad Sci U S A 68, 2112–2116. (doi:10.1073/pnas.68.9.2112)

9. Zehring WA, Wheeler DA, Reddy P, Konopka T-RJ, Kyriacou CP, Rosbash M, Hall JC. 1984 P-Element Transformation with period Locus DNA Restores Rhythmicity to Mutant, Arrhythmic Drosophila melanogaster. Cell. 39.

10. Bargiello TA, Jackson FR, Young MW. 1984 Restoration of circadian behavioural rhythms by gene transfer in Drosophila. Nature 312. (doi:10.1038/312752a0)

11. Vatine G, Vallone D, Gothilf Y, Foulkes NS. 2011 It’s time to swim! Zebrafish and the circadian clock. FEBS Lett 585, 1485–1494. (doi:10.1016/j.febslet.2011.04.007)

12. Tei H, Okamura H, Shigeyoshi Y, Fukuhara C, Ozawa R, Hirose M, Sakaki Y. 1997 Circadian oscillation of a mammalian homologue of the Drosophila period gene. Nature 389. (doi:10.1038/39086)

13. Takahashi JS. 2017 Transcriptional architecture of the mammalian circadian clock. Nat Rev Genet 18, 164–179. (doi:10.1038/nrg.2016.150)

14. van der Linden AM, Beverly M, Kadener S, Rodriguez J, Wasserman S, Rosbash M, Sengupta P. 2010 Genome-wide analysis of light- and temperature-entrained circadian transcripts in Caenorhabditis elegans. PLoS Biol 8, e1000503. (10.1016/j.cub.2004.11.057)

15. Olmedo M, O’Neill JS, Edgar RS, Valekunja UK, Reddy AB, Merrow M. 2012 Circadian regulation of olfaction and an evolutionarily conserved, nontranscriptional marker in Caenorhabditis elegans. Proc Natl Acad Sci U S A 109, 20479–20484. (doi:10.1073/pnas.1211705109)

16. Goya ME, Romanowski A, Caldart CS, Bénard CY, Golombek DA. 2016 Circadian rhythms identified in Caenorhabditis elegans by in vivo long-term monitoring of a bioluminescent reporter. Proc Natl Acad Sci U S A 113, E7837–E7845. (doi:10.1073/pnas.1605769113)

17. Simonetta SH, Golombek DA. 2007 An automated tracking system for Caenorhabditis elegans locomotor behavior and circadian studies application. J Neurosci Methods 161, 273–280. (doi:10.1016/j.jneumeth.2006.11.015)

18. Simonetta SH, Migliori ML, Romanowski A, Golombek DA. 2009 Timing of locomotor activity circadian rhythms in Caenorhabditis elegans. PLoS One 4, e7571. (doi:10.1371/journal.pone.0007571)

19. Herrero A, Romanowski A, Meelkop E, Caldart CS, Schoofs L, Golombek DA. 2015 Pigment-dispersing factor signaling in the circadian system of Caenorhabditis elegans. Genes Brain Behav 14, 493–501. (doi:10.1111/gbb.12231)

20. Saigusa T, Ishizaki S, Watabiki S, Ishii N, Tanakadate A, Tamai Y, Hasegawa K. 2002 Circadian behavioural rhythm in Caenorhabditis elegans [1]. Current Biology 12, 46–47. (doi:10.1016/S0960-9822(01)00669-8)

21. Winbush A, Gruner M, Hennig GW, van der Linden AM. 2015 Long-term imaging of circadian locomotor rhythms of a freely crawling C. elegans population. J Neurosci Methods 249, 66–74. (doi:10.1016/j.jneumeth.2015.04.009)

22. Caldart CS, Carpaneto A, Golombek DA. 2020 Synchronization of circadian locomotor activity behavior in Caernorhabditis elegans: Interactions between light and temperature. J Photochem Photobiol B 211, 112000. (doi:10.1016/j.jphotobiol.2020.112000)

23. Yu CW, Wu YC, Liao VHC. 2022 Early developmental nanoplastics exposure disturbs circadian rhythms associated with stress resistance decline and modulated by DAF-16 and PRDX-2 in C. elegans. J Hazard Mater 423. (doi:10.1016/j.jhazmat.2021.127091)

24. Migliori ML, Simonetta SH, Romanowski A, Golombek DA. 2011 Circadian rhythms in metabolic variables in Caenorhabditis elegans. Physiol Behav 103, 315–320. (doi:10.1016/j.physbeh.2011.01.026)

25. Migliori ML, Romanowski A, Simonetta SH, Valdez D, Guido M, Golombek DA. 2012 Daily variation in melatonin synthesis and arylalkylamine N-acetyltransferase activity in the nematode Caenorhabditis Elegans. J Pineal Res 53, 38–46. (doi:10.1111/j.1600-079X.2011.00969.x)

26. Kippert F, Saunders DS, Blaxter ML. 2002 Caenorhabditis elegans has a circadian clock [2]. Current Biology. 12. (doi:10.1016/S0960-9822(01)00670-4)

27. Hasegawa K, Saigusa T, Tamai Y. 2005 Caenorhabditis elegans opens up new insights into circadian clock mechanisms. Chronobiol Int 22, 1–19. (doi:10.1081/CBI-200038149)

28. Romanowski A, Garavaglia MJ, Goya ME, Ghiringhelli PD, Golombek DA. 2014 Potential conservation of circadian clock proteins in the phylum nematoda as revealed by bioinformatic searches. PLoS One 9, e112871. (doi:10.1371/journal.pone.0112871)

29. Hiroki S, Yoshitane H. 2024 Ror homolog nhr-23 is essential for both developmental clock and circadian clock in C. elegans. Commun Biol 7. (doi:10.1038/s42003-024-05894-3)

30. Marie A, Aguinaldo A, Turbevillet JM, Linford LS, Rivera MC, Garey, T: JR, Raff RA, Lake JA. 1997 Evidence for a clade of nematodes, arthropods and other moulting animals. Nature 387, 489–493.

31. Giribet G, Edgecombe GD. 2017 Current understanding of Ecdysozoa and its internal phylogenetic relationships. Integr Comp Biol 57, 455–466. (doi:10.1093/icb/icx072)

32. Moss EG. 2007 Heterochronic Genes and the Nature of Developmental Time. Current Biology. 17. (doi:10.1016/j.cub.2007.03.043)

33. Kim DH, Grün D, Van Oudenaarden A. 2013 Dampening of expression oscillations by synchronous regulation of a microRNA and its target. Nat Genet 45, 1337–1345. (doi:10.1038/ng.2763)

34. Hendriks GJ, Gaidatzis D, Aeschimann F, Großhans H. 2014 Extensive Oscillatory Gene Expression during C.elegans Larval Development. Mol Cell 53, 380–392. (doi:10.1016/j.molcel.2013.12.013)

35. Meeuse MW, Hauser YP, Morales Moya LJ, Hendriks G, Eglinger J, Bogaarts G, Tsiairis C, Großhans H. 2020 Developmental function and state transitions of a gene expression oscillator in Caenorhabditis elegans . Mol Syst Biol 16, 1–21. (doi:10.15252/msb.20209498)

36. Sun S, Rödelsperger C, Sommer RJ. 2021 Single worm transcriptomics identifies a developmental core network of oscillating genes with deep conservation across nematodes. Genome Res 31, 1590–1601. (doi:10.1101/gr.275303.121)

37. Filina O, Demirbas B, Haagmans R, van Zon JS. 2022 Temporal scaling in C. elegans larval development. Proc Natl Acad Sci U S A 119. (doi:10.1073/pnas.2123110119)

38. Meeuse MWM, Hauser YP, Nahar S, Smith AAT, Braun K, Azzi C, Rempfler M, Großhans H. 2023 C. elegans molting requires rhythmic accumulation of the Grainyhead/ LSF transcription factor GRH -1 . EMBO J 42. (doi:10.15252/embj.2022111895)

39. Olmedo M, Geibel M, Artal-Sanz M, Merrow M. 2015 A high-throughput method for the analysis of larval developmental phenotypes in Caenorhabditis elegans. Genetics 201, 443–448. (doi:10.1534/genetics.115.179242)

40. Jeon M, Gardner HF, Miller EA, Deshler J, Rougvie AE. 1999 Similarity of the C. elegans developmental timing protein LIN-42 to circadian rhythm proteins. Science (1979) 286. (doi:10.1126/science.286.5442.1141)

41. Bettinger JC, Lee K, Rougvie AE. 1996 Stage-specific accumulation of the terminal differentiation factor LIN-29 during Caenorhabditis elegans development. Development 122, 2517–2527. (doi:10.1242/dev.122.8.2517)

42. Banerjee D, Kwok A, Lin SY, Slack FJ. 2005 Developmental timing in C. elegans is regulated by kin-20 and tim-1, homologs of core circadian clock genes. Dev Cell 8, 287–295. (doi:10.1016/j.devcel.2004.12.006)

43. Tennessen JM, Gardner HF, Volk ML, Rougvie AE. 2006 Novel heterochronic functions of the Caenorhabditis elegans period-related protein LIN-42. Dev Biol 289, 30–43. (doi:10.1016/j.ydbio.2005.09.044)

44. Monsalve GC, Van Buskirk C, Frand AR. 2011 LIN-42/PERIOD controls cyclical and developmental progression of C. elegans molts. Current Biology 21, 2033–2045. (doi:10.1016/j.cub.2011.10.054)

45. Perales R, King DM, Aguirre-Chen C, Hammell CM. 2014 LIN-42, the Caenorhabditis elegans PERIOD homolog, Negatively Regulates MicroRNA Transcription. PLoS Genet 10. (doi:10.1371/journal.pgen.1004486)

46. Edelman TLB, McCulloch KA, Barr A, Frøkjær-Jensen C, Jorgensen EM, Rougvie AE. 2016 Analysis of a lin-42/period null allele implicates all three isoforms in regulation of caenorhabditis elegans molting and developmental timing. G3: Genes, Genomes, Genetics 6, 4077–4086. (doi:10.1534/g3.116.034165)

47. Rhodehouse K, Cascino K, Aseltine L, Padula A, Weinstein R, Spina JS, Olivero CE, Van Wynsberghe PM. 2018 The doubletime homolog KIN-20 mainly regulates let-7 independently of its effects on the period homolog LIN-42 in caenorhabditis elegans. G3: Genes, Genomes, Genetics 8, 2617–2629. (doi:10.1534/g3.118.200392)

48. Burr AH. 1985 THE PHOTOMOVEMENT OF Caenorhabditis elegans, A NEMATODE WHICH LACKS OCELLI. PROOF THAT THE RESPONSE IS TO LIGHT NOT RADIANT HEATING. Photochem Photobiol 41, 577–582. (doi:10.1111/j.1751-1097.1985.tb03529.x)

49. Ward A, Liu J, Feng Z, Xu XZS. 2008 Light-sensitive neurons and channels mediate phototaxis in C. elegans. Nat Neurosci 11, 916–922. (doi:10.1038/nn.2155)

50. Liu J et al. 2010 C. elegans phototransduction requires a G protein-dependent cGMP pathway and a taste receptor homolog. Nat Neurosci 13, 715–722. (doi:10.1038/nn.2540)

51. Gong J et al. 2016 The C. elegans Taste Receptor Homolog LITE-1 is a Photoreceptor. Cell 167, 1252–1263. (doi:10.1016/j.cell.2016.12.040)

52. Hanson SM et al. 2023 Structure-function analysis suggests that the photoreceptor LITE-1 is a light-activated ion channel. Current Biology 33. (doi:10.1016/j.cub.2023.07.008)

53. Haug MF, Gesemann M, Lazović V, Neuhauss SCF. 2015 Eumetazoan cryptochrome phylogeny and evolution. Genome Biol Evol 7, 601–619. (doi:10.1093/gbe/evv010)

54. Emery P, Venus So W, Kaneko M, Hall JC, Rosbash M. 1998 CRY, a Drosophila Clock and Light-Regulated Cryptochrome, Is a Major Contributor to Circadian Rhythm Resetting and Photosensitivity. Cell. 95.

55. Sato TK et al. 2006 Feedback repression is required for mammalian circadian clock function. Nat Genet 38, 312–319. (doi:10.1038/ng1745)

56. Ye R, Selby CP, Chiou YY, Ozkan-Dagliyan I, Gaddameedhi S, Sancar A. 2014 Dual modes of CLOCK:BMAL1 inhibition mediated by Cryptochrome and period proteins in the mammalian circadian clock. Genes Dev 28, 1989–1998. (doi:10.1101/gad.249417.114)

57. Petersen C, Krahn A, Leippe M. 2023 The nematode Caenorhabditis elegans and diverse potential invertebrate vectors predominantly interact opportunistically. Front Ecol Evol 11. (doi:10.3389/fevo.2023.1069056)

58. Barrière A, Félix MA. 2005 High local genetic diversity and low outcrossing rate in Caenorhabditis elegans natural populations. Current Biology 15, 1176– 1184. (doi:10.1016/j.cub.2005.06.022)

59. Kiontke KC, Félix MA, Ailion M, Rockman M V., Braendle C, Pénigault JB, Fitch DHA. 2011 A phylogeny and molecular barcodes for Caenorhabditis, with numerous new species from rotting fruits. BMC Evol Biol 11. (doi:10.1186/1471-2148-11-339)

60. Andersen EC, Gerke JP, Shapiro JA, Crissman JR, Ghosh R, Bloom JS, Félix MA, Kruglyak L. 2012 Chromosome-scale selective sweeps shape Caenorhabditis elegans genomic diversity. Nat Genet 44, 285–290. (doi:10.1038/ng.1050)

61. Sterken MG, Snoek LB, Kammenga JE, Andersen EC. 2015 The laboratory domestication of Caenorhabditis elegans. Trends in Genetics 31, 224–231. (doi:10.1016/j.tig.2015.02.009)

62. Schulenburg H, Félix MA. 2017 The natural biotic environment of Caenorhabditis elegans. Genetics 206. (doi:10.1534/genetics.116.195511)

63. Fielenbach N, Antebi A. 2008 C. elegans dauer formation and the molecular basis of plasticity. Genes Dev. 22, 2149–2165. (doi:10.1101/gad.1701508)

64. Hedgecock EM, Russell RL. 1975 Normal and mutant thermotaxis in the nematode Caenorhabditis elegans. Proc Natl Acad Sci U S A 72, 4061–4065. (doi:10.1073/pnas.72.10.4061)

65. Glauser DA. 2022 Temperature sensing and context-dependent thermal behavior in nematodes. Curr Opin Neurobiol. 73. (doi:10.1016/j.conb.2022.102525)

66. Lamberti ML et al. 2023 Regulation of the circadian clock in C. elegans by clock gene homologs kin-20 and lin-42. bioRxiv [Preprint] (doi:10.1101/2023.04.13.536481)

67. Glaser FT, Stanewsky R. 2005 Temperature synchronization of the Drosophila circadian clock. Current Biology 15, 1352–1363. (doi:10.1016/j.cub.2005.06.056)

68. Glaser FT, Stanewsky R. 2007 Synchronization of the Drosophila circadian clock by temperature cycles. Cold Spring Harb Symp Quant Biol 72, 233–242. (doi:10.1101/sqb.2007.72.046)

69. Harper REF, Ogueta M, Dayan P, Stanewsky R, Albert JT. 2017 Light Dominates Peripheral Circadian Oscillations in Drosophila melanogaster During Sensory Conflict. J Biol Rhythms 32, 423–432. (doi:10.1177/0748730417724250)

70. Lahiri K, Vallone D, Gondi SB, Santoriello C, Dickmeis T, Foulkes NS. 2005 Temperature regulates transcription in the zebrafish circadian clock. PLoS Biol 3, 2005–2016. (doi:10.1371/journal.pbio.0030351)

71. López-Olmeda JF, Madrid JA, Sánchez-Vázquez FJ. 2006 Light and temperature cycles as zeitgebers of zebrafish (Danio rerio) circadian activity rhythms. Chronobiol Int 23, 537–550. (doi:10.1080/07420520600651065)

72. Schurch NJ et al. 2016 How many biological replicates are needed in an RNA-seq experiment and which differential expression tool should you use? Rna 22, 839–851. (doi:10.1261/rna.053959.115)

73. Hughes ME, Hogenesch JB, Kornacker K. 2010 JTK-CYCLE: An efficient nonparametric algorithm for detecting rhythmic components in genome-scale data sets. J Biol Rhythms 25, 372–380. (doi:10.1177/0748730410379711)

74. Wu G, Anafi RC, Hughes ME, Kornacker K, Hogenesch JB. 2016 MetaCycle: An integrated R package to evaluate periodicity in large scale data. Bioinformatics 32, 3351–3353. (doi:10.1093/bioinformatics/btw405)

75. Langfelder P, Horvath S. 2008 WGCNA: An R package for weighted correlation network analysis. BMC Bioinformatics 9. (doi:10.1186/1471-2105-9-559)

76. Hughes ME, Hong HK, Chong JL, Indacochea AA, Lee SS, Han M, Takahashi JS, Hogenesch JB. 2012 Brain-specific rescue of Clock reveals system-driven transcriptional rhythms in peripheral tissue. PLoS Genet 8, e1002835. (doi:10.1371/journal.pgen.1002835)

77. Rund SSC, Gentile JE, Duffield GE. 2013 Extensive circadian and light regulation of the transcriptome in the malaria mosquito Anopheles gambiae. BMC Genomics 14. (doi:10.1186/1471-2164-14-218)

78. Hurley JM et al. 2014 Analysis of clock-regulated genes in Neurospora reveals widespread posttranscriptional control of metabolic potential. Proc Natl Acad Sci U S A 111, 16995–17002. (doi:10.1073/pnas.1418963111)

79. Yang Y, Li Y, Sancar A, Oztas O. 2020 The circadian clock shapes the Arabidopsis transcriptome by regulating alternative splicing and alternative polyadenylation. Journal of Biological Chemistry 295, 7608–7619. (doi:10.1074/jbc.RA120.013513)

80. Huang DW, Sherman BT, Lempicki RA. 2009 Systematic and integrative analysis of large gene lists using DAVID bioinformatics resources. Nat Protoc 4, 44–57. (doi:10.1038/nprot.2008.211)

81. Huang DW, Sherman BT, Lempicki RA. 2009 Bioinformatics enrichment tools: Paths toward the comprehensive functional analysis of large gene lists. Nucleic Acids Res 37, 1–13. (doi:10.1093/nar/gkn923)

82. Janssen T, Husson SJ, Lindemans M, Mertens I, Rademakers S, Ver Donck K, Geysen J, Jansen G, Schoofs L. 2008 Functional characterization of three G protein-coupled receptors for pigment dispersing factors in Caenorhabditis elegans. Journal of Biological Chemistry 283, 15241–15249. (doi:10.1074/jbc.M709060200)

83. Yoo SH et al. 2004 PERIOD2::LUCIFERASE real-time reporting of circadian dynamics reveals persistent circadian oscillations in mouse peripheral tissues. Proc Natl Acad Sci U S A 101, 5339–5346. (doi:10.1073/pnas.0308709101)

84. Welsh DK, Yoo S-H, Liu AC, Takahashi JS, Kay SA. 2004 Bioluminescence Imaging of Individual Fibroblasts Reveals Persistent, Independently Phased Circadian Rhythms of Clock Gene Expression. Current Biology 14, 2289– 2295. (doi:10.1016/j.cub.2004.11.057)

85. Keegan KP, Pradhan S, Wang JP, Allada R. 2007 Meta-analysis of Drosophila circadian microarray studies identifies a novel set of rhythmically expressed genes. PLoS Comput Biol 3, 2087–2110. (doi:10.1371/journal.pcbi.0030208)

86. Brooks TG, Manjrekar A, Mrcěla A, Grant GR. 2023 Meta-analysis of Diurnal Transcriptomics in Mouse Liver Reveals Low Repeatability of Rhythm Analyses. J Biol Rhythms 38, 556–570. (doi:10.1177/07487304231179600)

87. Mure LS et al. 2018 Diurnal transcriptome atlas of a primate across major neural and peripheral tissues. Science (1979) 359. (doi:10.1126/science.aao0318)

88. Mei W, Jiang Z, Chen Y, Chen L, Sancar A, Jiang Y. 2021 Genome-wide circadian rhythm detection methods: Systematic evaluations and practical guidelines. Brief Bioinform. 22. (doi:10.1093/bib/bbaa135)

89. Goya ME, Romanowski A, Caldart CS, Bénard CY, Golombek DA. 2016 Circadian rhythms identified in Caenorhabditis elegans by in vivo long-term monitoring of a bioluminescent reporter. Proc Natl Acad Sci U S A 113, E7837–E7845. (doi:10.1073/pnas.1605769113)

90. Pan Y et al. 2020 12-h clock regulation of genetic information flow by XBP1s. PLoS Biol 18. (doi:10.1371/journal.pbio.3000580)

91. Meng H, Gonzales NM, Lonard DM, Putluri N, Zhu B, Dacso CC, York B, O’Malley BW. 2020 XBP1 links the 12-hour clock to NAFLD and regulation of membrane fluidity and lipid homeostasis. Nat Commun 11. (doi:10.1038/s41467-020-20028-z)

92. Zhu et al. 2023 Evidence for conservation of primordial ∼12-hour ultradian gene programs in humans under free-living conditions. bioRxiv [preprint] (doi:10.1101/2023.05.02.539021)

93. Zhu B, Zhang Q, Pan Y, Mace EM, York B, Antoulas AC, Dacso CC, O’Malley BW. 2017 A Cell-Autonomous Mammalian 12 hr Clock Coordinates Metabolic and Stress Rhythms. Cell Metab 25, 1305–1319.e9. (doi:10.1016/j.cmet.2017.05.004)

94. Pembroke WG, Babbs A, Davies KE, Ponting CP, Oliver PL. 2015 Temporal transcriptomics suggest that twin-peaking genes reset the clock. Elife 4, 1–15. (doi:10.7554/elife.10518)

95. Ye H et al. 2021 Gene co-expression analysis identifies modules related to insufficient sleep in humans. Sleep Med 86, 68–74. (doi:10.1016/j.sleep.2021.08.010)

96. Poggiogalle E, Jamshed H, Peterson CM. 2018 Circadian regulation of glucose, lipid, and energy metabolism in humans. Metabolism 84. (doi:10.1016/j.metabol.2017.11.017)

97. McCarthy JJ et al. 2007 Identification of the circadian transcriptome in adult mouse skeletal muscle. Physiol Genomics 31. (doi:10.1152/physiolgenomics.00066.2007)

98. Jouffe C, Cretenet G, Symul L, Martin E, Atger F, Naef F, Gachon F. 2013 The Circadian Clock Coordinates Ribosome Biogenesis. PLoS Biol 11. (doi:10.1371/journal.pbio.1001455)

99. An H, Harper JW. 2020 Ribosome Abundance Control Via the Ubiquitin– Proteasome System and Autophagy. J Mol Biol. 432. (doi:10.1016/j.jmb.2019.06.001)

100. Nagaraj N, Wisniewski JR, Geiger T, Cox J, Kircher M, Kelso J, Pääbo S, Mann M. 2011 Deep proteome and transcriptome mapping of a human cancer cell line. Mol Syst Biol 7. (doi:10.1038/msb.2011.81)

101. Wiśniewski JR, Hein MY, Cox J, Mann M. 2014 A ‘proteomic ruler’ for protein copy number and concentration estimation without spike-in standards. Molecular and Cellular Proteomics 13. (doi:10.1074/mcp.M113.037309)

102. Seinkmane E et al. 2024 Circadian regulation of macromolecular complex turnover and proteome renewal. EMBO J (doi:10.1038/s44318-024-00121-5)

103. Dörner K, Ruggeri C, Zemp I, Kutay U. 2023 Ribosome biogenesis factors— from names to functions. EMBO J 42. (doi:10.15252/embj.2022112699)

104. Hughes ME, DiTacchio L, Hayes KR, Vollmers C, Pulivarthy S, Baggs JE, Panda S, Hogenesch JB. 2009 Harmonics of circadian gene transcription in mammals. PLoS Genet 5. (doi:10.1371/journal.pgen.1000442)

105. Yoshitane H et al. 2014 CLOCK-Controlled Polyphonic Regulation of Circadian Rhythms through Canonical and Noncanonical E-Boxes. Mol Cell Biol 34, 1776–1787. (doi:10.1128/mcb.01465-13)

106. Correa A, Lewis ZA, Greene A V., March IJ, Gomer RH, Bell-Pedersen D. 2003 Multiple oscillators regulate circadian gene expression in Neurospora. Proc Natl Acad Sci U S A 100. (doi:10.1073/pnas.2233734100)

107. Rodriguez J, Tang CHA, Khodor YL, Vodala S, Menet JS, Rosbash M. 2013 Nascent-Seq analysis of Drosophila cycling gene expression. Proc Natl Acad Sci U S A 110, E275–84. (doi:10.1073/pnas.1219969110)

108. Nakajima M, Imai K, Ito H, Nishiwaki T, Murayama Y, Iwasaki H, Oyama T, Kondo T. 2005 Reconstitution of circadian oscillation of cyanobacterial KaiC phosphorylation in vitro. Science (1979) 308, 414–415. (doi:10.1126/science.1108451)

109. O’Neill JS, Reddy AB. 2011 Circadian clocks in human red blood cells. Nature 469, 498–504. (doi:10.1038/nature09702)

110. Ray S, Valekunja UK, Stangherlin A, Howell SA, Snijders AP, Damodaran G, Reddy AB. In press. Circadian rhythms in the absence of the clock gene Bmal1.

111. Lipton JO et al. 2015 The circadian protein BMAL1 regulates translation in response to S6K1-mediated phosphorylation. Cell 161. (doi:10.1016/j.cell.2015.04.002)

112. Wong DCS et al. 2022 CRYPTOCHROMES promote daily protein homeostasis. EMBO J 41, 1–23. (doi:10.15252/embj.2021108883)

113. Putker M et al. 2021 CRYPTOCHROMES confer robustness, not rhythmicity, to circadian timekeeping. EMBO J 40, 1–15. (doi:10.15252/embj.2020106745)

114. Kwiatkowski ER, Schnytzer Y, Rosenthal JJC, Emery P. 2023 Behavioral circatidal rhythms require Bmal1 in Parhyale hawaiensis. Current Biology 33. (doi:10.1016/j.cub.2023.03.015)

115. Lin Z, Green EW, Webster SG, Hastings MH, Wilcockson DC, Kyriacou CP. 2023 The circadian clock gene bmal1 is necessary for co-ordinated circatidal rhythms in the marine isopod Eurydice pulchra (Leach). PLoS Genet 19. (doi:10.1371/journal.pgen.1011011)

116. O’Neill JS, Lee KD, Zhang L, Feeney K, Webster SG, Blades MJ, Kyriacou CP, Hastings MH, Wilcockson DC. 2015 Metabolic molecular markers of the tidal clock in the marine crustacean Eurydice pulchra. Current Biology. 25. (doi:10.1016/j.cub.2015.02.052)

117. Zhang L, Hastings MH, Green EW, Tauber E, Sladek M, Webster SG, Kyriacou CP, Wilcockson DC. 2013 Dissociation of circadian and circatidal timekeeping in the marine crustacean eurydice pulchra. Current Biology 23. (doi:10.1016/j.cub.2013.08.038)

118. Tran D, Perrigault M, Ciret P, Payton L. 2020 Bivalve mollusc circadian clock genes can run at tidal frequency. Proceedings of the Royal Society B: Biological Sciences 287. (doi:10.1098/rspb.2019.2440)

119. Chawla S et al. 2024 Timely Questions Emerging in Chronobiology: The Circadian Clock Keeps on Ticking. J Circadian Rhythms. 22. (doi:10.5334/jcr.237)

120. Thomas JH. 1990 Genetic Analysis of Defecation in Caenorhabditis elegans. Genetis 124, 855–892.

121. Kaulich E, Carroll T, Ackley BD, Tang YQ, Hardege I, Nehrke K, Schafer WR, Walker DS. 2022 Distinct roles for two Caenorhabditis elegans acid-sensing ion channels in an ultradian clock. Elife 11. (doi:10.7554/eLife.75837)

122. Collins plus some more. In press. Activity of the C. elegans egg-laying behavior circuit is controlled by competing activation and feedback inhibition. (doi:10.7554/eLife.21126.001)

123. Kinney B et al. 2023 A circadian-like gene network programs the timing and dosage of heterochronic miRNA transcription during C. elegans development. Dev Cell 58. (doi:10.1016/j.devcel.2023.08.006)

124. MR K, DI H. 1981 Sperm isolation and biochemical analysis of the major sperm protein from C. elegans. Dev Biol 84, 299–312.

125. Roberts TM, Ward S. 1982 Centripetal flow of pseudopodial surface components could propel the amoeboid movement of Caenorhabditis elegans spermatozoa. Journal of Cell Biology 92, 132–138. (doi:10.1083/jcb.92.1.132)

126. Miller MA, Nguyen VQ, Lee MH, Kosinski M, Schedl T, Caprioli RM, Greenstein D. 2001 A sperm cytoskeletal protein that signals oocyte meiotic maturation and ovulation. Science (1979) 291, 2144–2147. (doi:10.1126/science.1057586)

127. Huelgas-Morales G, Greenstein D. 2018 Control of oocyte meiotic maturation in C. elegans. Semin Cell Dev Biol. 84, 90–99. (doi:10.1016/j.semcdb.2017.12.005)

128. Stiernagle T. 2006 Maintenance of C. elegans. WormBook. (doi:10.1895/wormbook.1.101.1)

129. Mitchell DH, Stiles JW, Santelli J, Rao Sanadi D. 1979 Synchronous growth and aging of Caenorhabditis elegans in the presence of fluorodeoxyuridine. Journals of Gerontology 34, 28–36. (doi:10.1093/geronj/34.1.28)

130. Ewels P, Magnusson M, Lundin S, Käller M. 2016 MultiQC: Summarize analysis results for multiple tools and samples in a single report. Bioinformatics 32, 3047–3048. (doi:10.1093/bioinformatics/btw354)

131. Martin M. 2011 Cutadapt removes adapter sequences from high-throughput sequencing reads. EMBnet J 17, 10. (doi:10.14806/ej.17.1.200)

132. Patro R, Duggal G, Love MI, Irizarry RA, Kingsford C. 2017 Salmon provides fast and bias-aware quantification of transcript expression. Nat Methods 14, 417–419. (doi:10.1038/nmeth.4197)

133. Benson DA, Cavanaugh M, Clark K, Karsch-Mizrachi I, Lipman DJ, Ostell J, Sayers EW. 2013 GenBank. Nucleic Acids Res 41, 36–42. (doi:10.1093/nar/gks1195)

134. Pimentel H, Bray NL, Puente S, Melsted P, Pachter L. 2017 Differential analysis of RNA-seq incorporating quantification uncertainty. Nat Methods 14, 687–690. (doi:10.1038/nmeth.4324)

135. Team RC. 2023 R Core Team 2023 R: A language and environment for statistical computing. R foundation for statistical computing. https://www.R-project.org/. R Foundation for Statistical Computing

136. Posit team. 2023 RStudio: Integrated Development for R. RStudio, Inc., Boston, MA.

137. Johnson WE, Li C, Rabinovic A. 2007 Adjusting batch effects in microarray expression data using empirical Bayes methods. Biostatistics 8, 118–127. (doi:10.1093/biostatistics/kxj037)

138. Leek JT, Johnson WE, Parker HS, Jaffe AE, Storey JD. 2012 The SVA package for removing batch effects and other unwanted variation in high-throughput experiments. Bioinformatics 28. (doi:10.1093/bioinformatics/bts034)

139. Wickham H. 2016 ggplot2 Elegant Graphics for Data Analysis (Use R!). Springer

