## Supplementary figures for "The rhythmic transcriptional landscape in *Caenorhabditis elegans*: daily, circadian and novel 16-hour cycling gene expression revealed by RNA-sequencing"

Figure S1

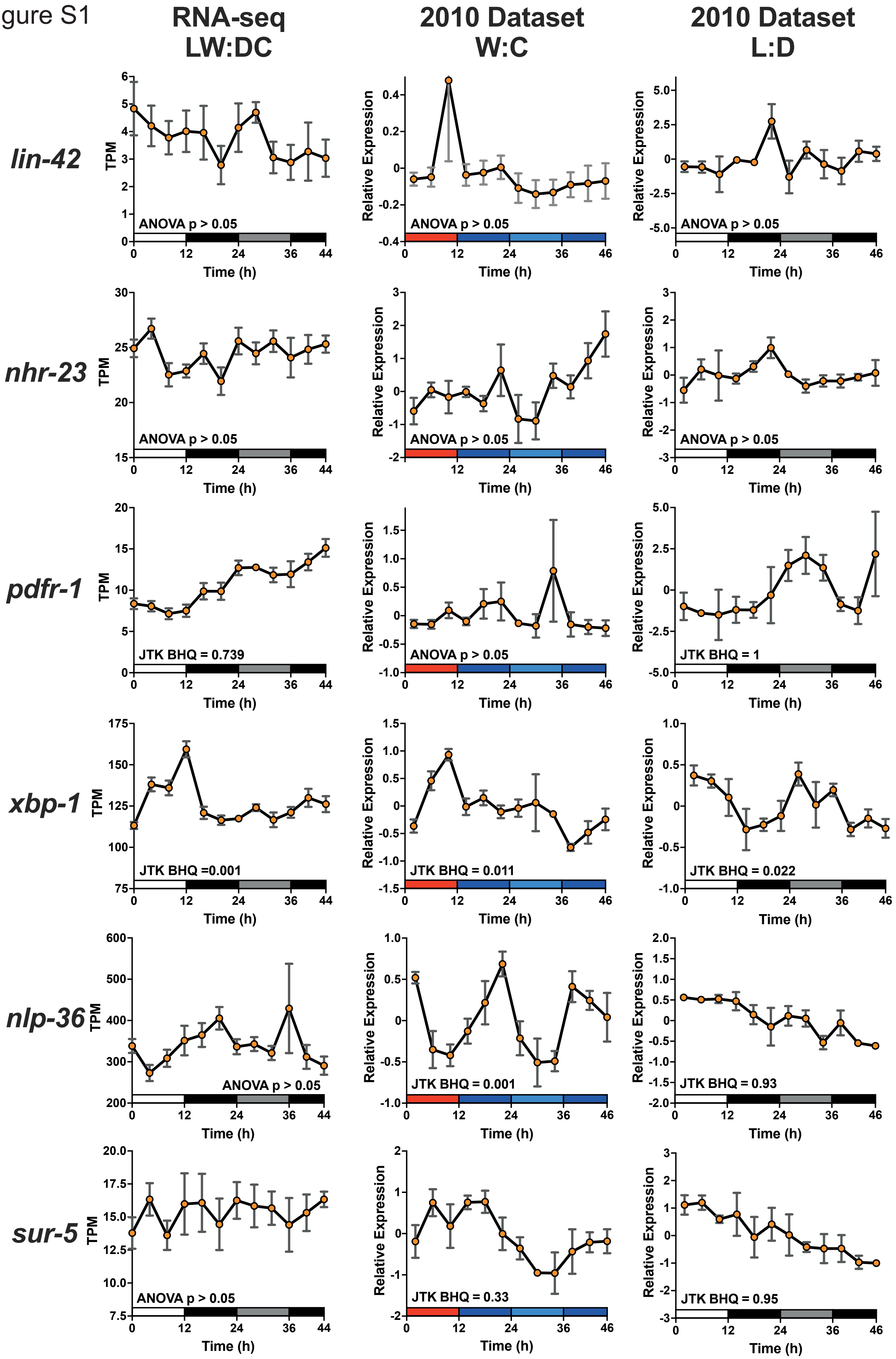

RNA-seq
LW:DC
2010 Dataset
W:C
2010 Dataset
L:D
lin-42
nhr-23
pdfr-1
xbp-1
nlp-36
sur-5
TPM
Relative Expression
Time (h)
ANOVA p > 0.05
JTK BHQ = 0.739
ANOVA p > 0.05
JTK BHQ = 1
JTK BHQ =0.001
JTK BHQ = 0.011
JTK BHQ = 0.022
ANOVA p > 0.05
JTK BHQ = 0.001
JTK BHQ = 0.93
ANOVA p > 0.05
JTK BHQ = 0.33
JTK BHQ = 0.95

**Supplementary Figure 1: Expression profiles comparing specific genes of interest across three independent datasets.** Shown are average expression values of replicates ± SEM. JTK_Cycle BHQ value presented where ANOVA $p < 0.05$, ANOVA $p$-value presented otherwise. For the 2010 dataset, relative expression was calculated previously as described [14]. Time 0 represents the time of light onset and/or temperature increases respectively with environmental conditions indicated by boxes along the $x$-axis (**left column: RNA-seq,** white = light/20°C, black = dark/15°C, light grey = subjective day (dark/15°C); **Centre: 2010 dataset temperature entrainment,** red = 25°C, blue = 15°C, light blue = subjective day (15°C)). **Right: 2010 microarray dataset light entrainment,** white = light, black = dark, grey = subjective day (dark), all at 18°C.

Figure S2

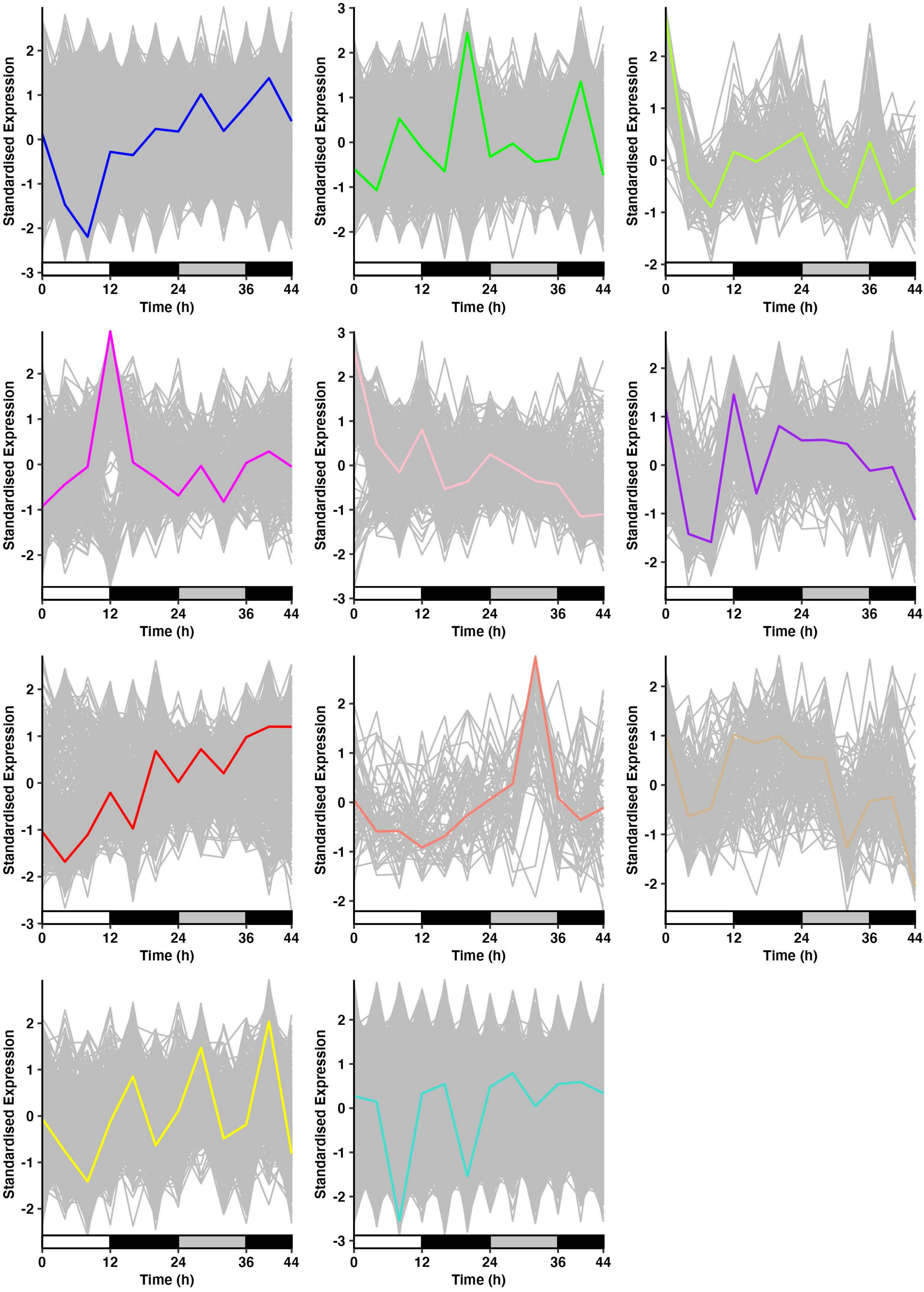

**Supplementary Figure 2: Additional WGNCA modules.** Shown are all 11 remaining modules identified by WGCNA not presented in the main text. Coloured line indicates the module designation and the eigengene expression averaged by time point. Modules are (from left to right): Blue, Green, Greenyellow, Magenta, Pink, Purple, Red, Salmon, Tan, Yellow and Turquoise. Bars along $x$-axis indicate environmental conditions: white = light/20°C, dark grey = dark/15°C, light grey = subjective day (dark/15°C)). Genes not assigned to a module (3436) are not shown. Full gene assignments given in Table S8.
